# Proteome-wide copy-number estimation from transcriptomics

**DOI:** 10.1101/2023.07.10.548432

**Authors:** Andrew J. Sweatt, Cameron D. Griffiths, B. Bishal Paudel, Kevin A. Janes

## Abstract

Protein copy numbers constrain systems-level properties of regulatory networks, but absolute proteomic data remain scarce compared to transcriptomics obtained by RNA sequencing. We addressed this persistent gap by relating mRNA to protein statistically using best-available data from quantitative proteomics–transcriptomics for 4366 genes in 369 cell lines. The approach starts with a central estimate of protein copy number and hierarchically appends mRNA-protein and mRNA-mRNA dependencies to define an optimal gene-specific model that links mRNAs to protein. For dozens of independent cell lines and primary prostate samples, these protein inferences from mRNA outmatch stringent null models, a count-based protein-abundance repository, and empirical protein-to-mRNA ratios. The optimal mRNA-to-protein relationships capture biological processes along with hundreds of known protein-protein interaction complexes, suggesting mechanistic relationships are embedded. We use the method to estimate viral-receptor abundances of CD55–CXADR from human heart transcriptomes and build 1489 systems-biology models of coxsackievirus B3 infection susceptibility. When applied to 796 RNA sequencing profiles of breast cancer from The Cancer Genome Atlas, inferred copy-number estimates collectively reclassify 26% of Luminal A and 29% of Luminal B tumors. Protein-based reassignments strongly involve a pharmacologic target for luminal breast cancer (CDK4) and an α-catenin that is often undetectable at the mRNA level (CTTNA2). Thus, by adopting a gene-centered perspective of mRNA-protein covariation across different biological contexts, we achieve accuracies comparable to the technical reproducibility limits of contemporary proteomics. The collection of gene-specific models is assembled as a web tool for users seeking mRNA-guided predictions of absolute protein abundance (http://janeslab.shinyapps.io/Pinferna).

**Significance statement:** It is easier to quantify mRNA in cells than it is to quantify protein, but proteins are what execute most cellular functions. Even though protein is synthesized from mRNA in cells, relating a cellular quantity of mRNA to a quantity of protein is challenging. Here, we bring together quantitative measures of mRNA and protein for 4366 genes in 369 different cultured cell types to build equations that predict protein abundance from the abundance of mRNAs expressed. These equations capture facets of biological regulation and work better than existing alternatives that rely on consensus values or ratios. Since mRNA measurements are more widespread than protein, this study makes new analyses possible by protein estimation from mRNA.

## Introduction

Absolute numbers of molecules place important bounds on biological systems, but they are hard to come by (1). One exception is deep RNA sequencing (RNA-seq) of bulk samples, which provides absolute transcript-per-million (TPM) estimates of all expressed genes (2). The commoditization of sequencing has made RNA-seq the prevailing omics approach: as of mid-2023, the leading repository (3) contains >32,000 studies with human samples. RNA-seq profiles are useful for reading out the state of the genome (4, 5), but mapping the transcriptome to the abundance of proteins is complex. In tumor classification, for example, the number and identity of cancer subtypes changes when using quantitative measurements of mRNA versus protein (6, 7). The challenge is especially acute for mathematical models in systems biology, which need protein quantities to constrain topology (8), initial conditions (9), or transition rates (10). Filling the overall gap requires new strategies for absolute quantification of proteomes for different needs.

Progressive experimental innovations in untargeted mass spectrometry have made quantitative proteomics a reality (11). Isobaric labeling approaches such as tandem mass tagging (TMT) (12) now quantify 11+ multiplex samples and are the method of choice for proteogenomics (13). Comparisons of individual proteins across samples indicate that linear mRNA–protein relationships vary greatly in quality (Pearson *R* = –0.4 to 0.8) depending on the gene and gene category (6, 7). Unfortunately, multiplex labeling yields peptide-specific relative quantities that cannot examine absolute differences among proteins within a sample (11). This difficulty is surmounted by data-independent acquisition methods like sequential window acquisition of all theoretical mass spectra (SWATH) (14), which analyzes all precursor ions in a series of mass-to-charge ratio windows. After data acquisition, the most sensitive peptide(s) of a protein are summed by intensity, and the resulting data are centered at a reasonable per-cell average (10^4^ copies) to yield absolute estimates of the detectable proteome. SWATH is newer, harder to adapt to different cell lineages, and lower throughput. Consequently, the leading proteomics repository (15) contains ∼tenfold fewer SWATH depositions than label-based depositions as of mid-2023. The need for commoditized SWATH-like protein estimates may forever outpace the ability to generate them directly.

As a means for bootstrapping absolute protein copy numbers, an appealing starting point is RNA-seq. mRNA is the template for protein translation, and in terms of scale, depositions of human RNA-seq exceed those of quantitative proteomics (all species, all methods) by ∼fivefold (3, 15). However, despite useful transcriptomic inference of protein activities from the gene networks surrounding them (16), directly estimating absolute protein copy numbers from mRNA is historically fraught with uncertainty. Linear mRNA–protein relationships adequately recapitulate protein expression among genes within a sample, but they are poor at distinguishing protein differences among samples for any given gene (17–20). The latter is important for systems biology when using transcriptome profiles to instantiate personalized models of function (8–10). The current thinking is that the steady-state abundance of mRNA and its intrinsic translation rate create a general “set point” for protein expression, which is buffered or tuned according to the abundance of complexes that stably contain the protein (19, 21). Unfortunately, our working inventory of protein-protein interactions and stable complexes in mammalian cells is far from saturation (22, 23), which has thus far prevented a bottom-up reconstruction of mRNA-to-protein relationships that are absolute and conditional.

Here, we surmounted this challenge by adopting a top-down perspective that statistically identifies the best working absolute mRNA-to-protein relationship for each gene based on paired data in several hundred cancer cell lines. SWATH and TMT datasets from different sources are encouragingly self-consistent, enabling the meta-assembly used for model training and selection of three relationship classes. Although relationship classes are entirely data driven, we find biological meaning and gene-specific mechanisms in each. The approach consistently improves the accuracy of proteome-wide inferences from RNA-seq transcriptomes of cells, tumors, and tissues when compared to other tools and a stringent null hypothesis specific to each gene’s protein set point. We use the method to build 1489 personalized systems-biology models of virus infection (24, 25) and re-classify 796 cases of breast cancer (26) according to inferred absolute protein abundance from public RNA-seq datasets. This study provides an open and accessible route to gleaning protein copy numbers from RNA-seq when it is impractical or impossible to quantify the proteome directly (http://janeslab.shinyapps.io/Pinferna).

## Results

### Deriving three gene-specific biological classes of mRNA-protein relationships

To estimate mRNA-protein relationships, we obtained quantitative proteomics measured by TMT mass spectrometry in 375 cancer cell lines (27) and placed these data on an absolute scale by using independent SWATH proteomics from two lines—one breast carcinoma (28) and one osteosarcoma (29)—within the TMT dataset (Fig. 1*A*, Step 1, and *SI Appendix,* Fig. S1*A* and Table S1). Training with a large, diverse panel of cancer lines avoids confounding gene or protein covariations that may arise in primary tissues and tumors because of cell mixtures (30). When TMT profiles scaled to one reference line were compared to SWATH data measured directly in the other reference line, correlations were above 0.7 in both cases (*p* < 10^-15^; Fig. 1*B* and *SI Appendix*, *F*ig. S1*B*), placing bounds on the internal consistency of the two data sources. Overall, the meta-assembly yielded protein copy number measurements for 4384 proteins across 375 cell lines.

**Fig. 1.**
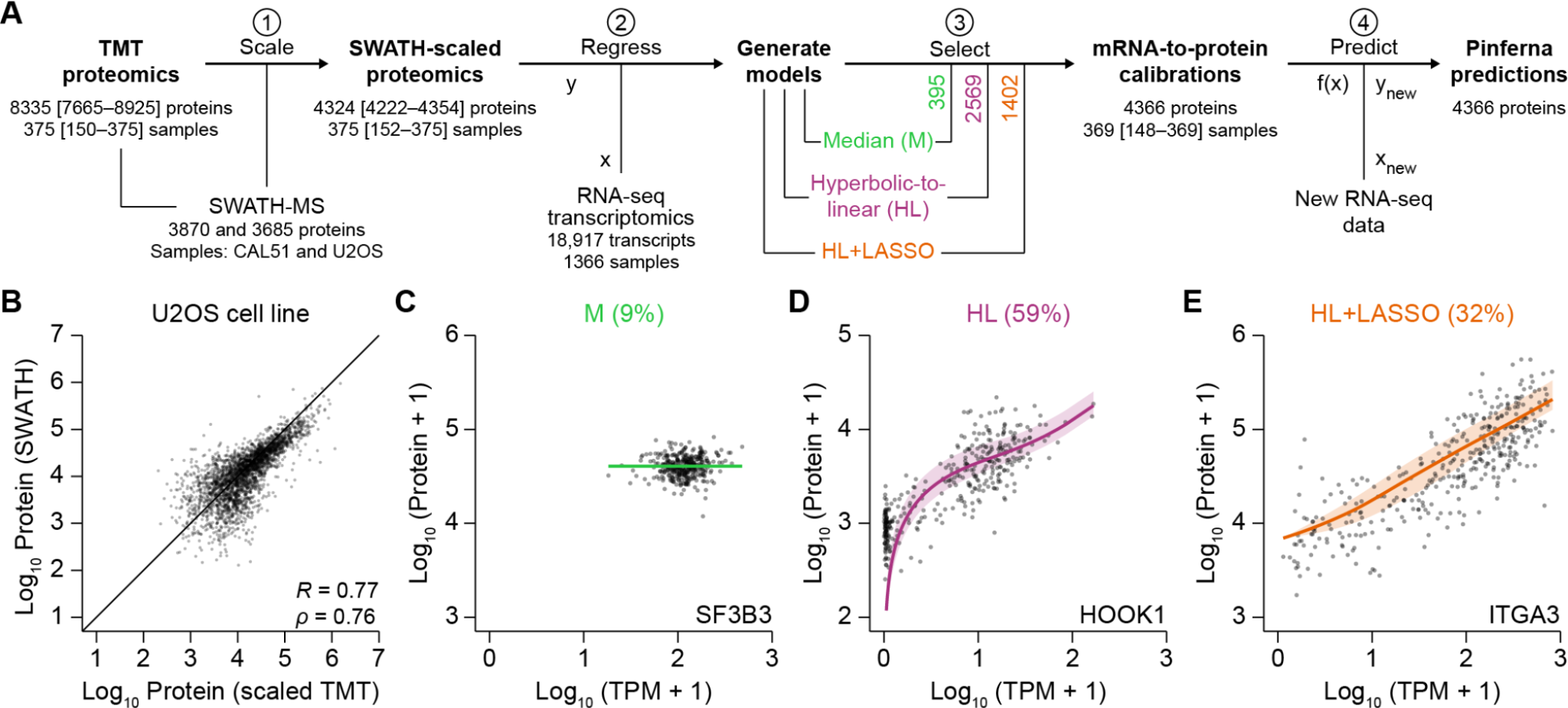
Meta-assembly and inference of conditional mRNA-to-protein relationships for 4366 human genes. (*A*) Data fusion and model discrimination. (1) Tandem mass tag (TMT) proteomics of 375 cancer cell lines (27) were calibrated to an absolute scale based on sequential window acquisition of all theoretical mass spectra (SWATH) proteomics of CAL51 and U2OS cells (PXD003278; PXD000954). (2) SWATH-scaled proteins were regressed using three models that incorporate transcript abundance from RNA sequencing (RNA-seq) to different extents: median (M), no contribution of mRNA; hyperbolic-to-linear (HL) relationship incorporating mRNA of the gene, 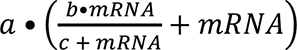; HL + least absolute shrinkage and selection operator (LASSO) regressors with mRNAs other than the gene of interest. (3) Model selection for each gene was based on the Bayesian Information Criterion. The number of genes selected in each class is indicated. (4) New samples profiled by RNA-seq were used with the calibrated models to make protein inference from RNA (Pinferna) predictions. The number of proteins measured per sample or number of samples with data per protein is shown at each step as the median with the range in brackets. (*B*) Reliable cross-calibration of the TMT and SWATH meta-assembly. Step 1 of Fig. 1A was performed with CAL51 data alone and the SWATH-scaled TMT proteomics of U2OS cells compared with data obtained directly by SWATH. Pearson’s *R* and Spearman’s *ρ* are shown. The reciprocal cross-calibration is shown in *SI Appendix*, Fig. S1*B*. (*C–E*) Representative M, HL, and HL+LASSO genes. Absolute protein copies per cell were regressed against the mRNA abundance normalized as transcripts per million (TPM). Best-fit calibrations ± 95% confidence intervals are overlaid on the proteomic–transcriptomic data from *n* = 369 cancer cell lines. Evidence for model selection is shown in *SI Appendix,* Fig. S1*C–E*.

We discerned mathematical relationships that best captured abundance relationships between mRNA and protein by merging SWATH-scaled proteomics with paired Cancer Cell Line Encyclopedia transcriptomics (31) and building gene-specific regressions of different classes (Fig. 1*A*, Step 2, and *SI Appendix*, Tables S2–S4). In the simplest case, protein abundance varies nominally around the median (M) regardless of transcript abundance (Fig. 1*C*). To incorporate abundance information about the transcript, we also evaluated a hyperbolic-to-linear (HL) relationship where low-abundance changes in mRNA cause larger nonlinear changes in protein that linearize as mRNA abundance increases (Fig. 1*D*). Finally, we considered that the abundance of some proteins may be further explained by the abundance of other transcripts and applied the least absolute shrinkage and selection operator (LASSO) to residuals of the HL fit. The best model among the three for each gene was distinguished by the Bayesian Information Criterion (BIC; Fig. 1*A*, step 3, and *SI Appendix,* Fig. S1*C–E*) to arrive at preferred M, HL, or HL+LASSO relationships for 4366 genes (*SI Appendix*, Table S5). The best model for each gene was strongly preferred over the others in 98% of cases (*SI Appendix*, Fig. S1*F*). These relationships create a template for <u>p</u>rotein <u>infer</u>ence from <u>RNA</u> (Pinferna) given new samples with transcriptomic profiles (Fig. 1*A*, step 4).

We examined characteristics of the genes in each model class. Consistent with previous findings (6, 32), M genes with no clear transcript dependence showed gene ontology (GO) enrichments for translation and mitochondrial electron transport (Fig. 2*A* and *SI Appendix,* Table S6). Although M genes were not significantly longer lived than others (*SI Appendix,* Fig. S2*A,B*), we found that they had high transcript abundances (*SI Appendix,* Fig. S2*C*) and were enriched in multi-protein complexes in the CORUM database (*p* < 10^-59^) (33). Proteins residing in stable complexes may saturate for all measured abundances of mRNA because their copy numbers are stoichiometrically limited by other subunits (21), causing the loss of an observable relationship between mRNA and protein.

**Fig. 2.**
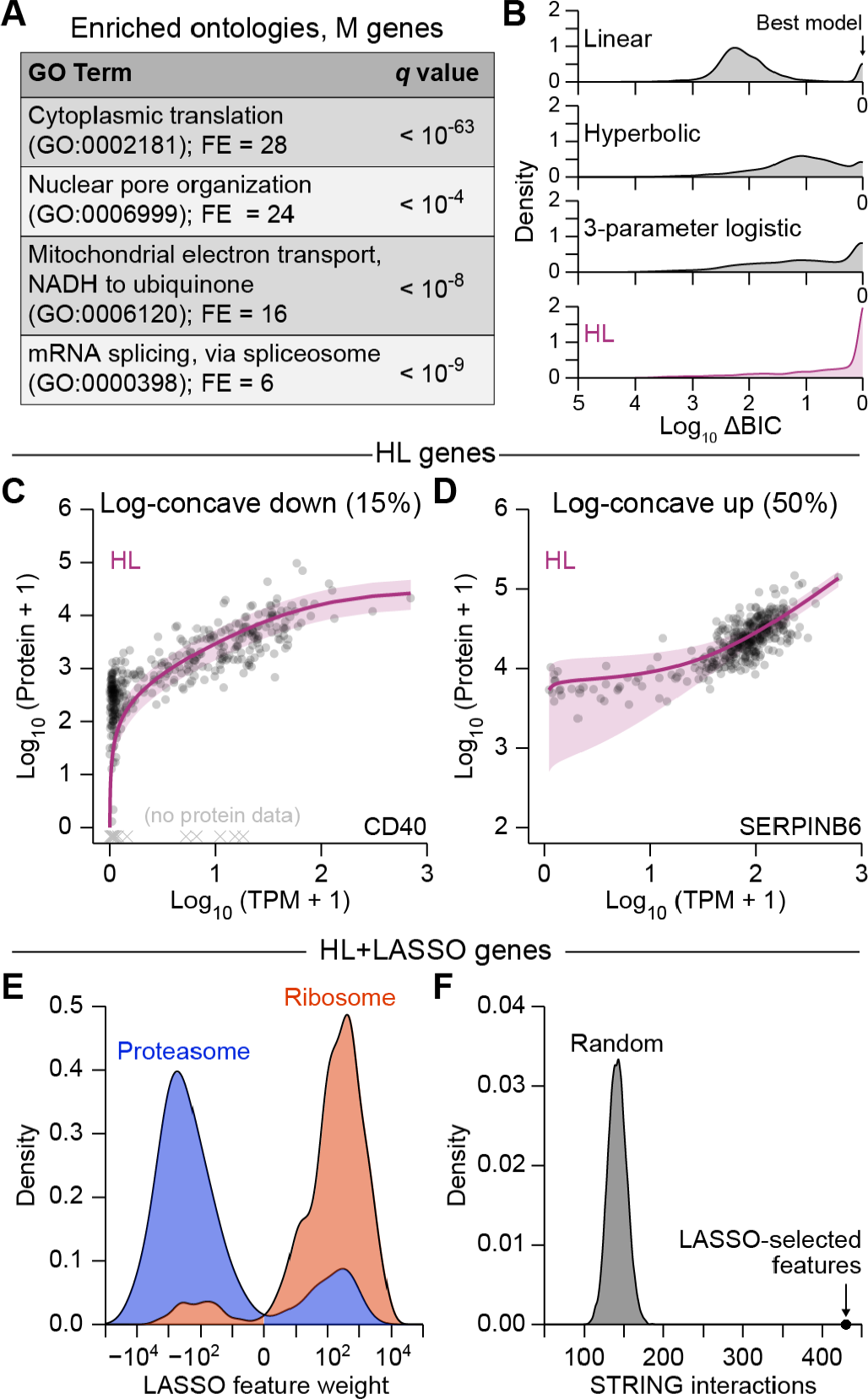
Pinferna model selection is consistent with known biological mechanisms and mRNA-to-protein relationships. (*A*) Gene ontology (GO) enrichments for M genes. The largest non-redundant GO term is shown with the fold enrichment (FE) and false discovery rate-corrected *p* value (*q*). The complete list of GO enrichments for each relationship class is available in *SI Appendix,* Table S6. (*B*) HL outperforms competing mRNA-to-protein relationships. Models encoding linear, hyperbolic, three-parameter logistic, and HL relationships were built for all genes (*n* = 4366) and compared by Bayesian Information Criterion (BIC). Results are shown as the smoothed density of BIC differences (ΔBIC) relative to the best model for that gene (ΔBIC = 0). Distributions of BIC weights (34) are shown in *SI Appendix,* Fig. S2*D*. (*C**–**D*) HL captures different empirical classes of mRNA-to-protein relationships. Log concave-down genes (*C*) saturate at high mRNA abundance, whereas log concave-up genes (*D*) plateau at low mRNA abundance. The remaining genes exhibited characteristics of both fits or linear relationships to varying degrees (*SI Appendix,* Fig. S2*G–I*). (*E*) Feature weights of HL+LASSO genes are biologically sensible. Smoothed densities of LASSO feature weights (indicating strength and direction of modulation for an HL fit) among mRNAs encoding subunits of the proteasome (*n* = 127 feature weights; blue) and the ribosome (*n* = 397 feature weights; red) are shown. (*F*) HL+LASSO feature weights are highly enriched for STRING interactions. For each HL+LASSO gene, LASSO-selected features were replaced with random genes (*n* = 10,000 iterations) to build a null distribution for finding binary interactions in STRING (36). The actual number of STRING interactions among HL+LASSO genes of Pinferna is indicated.

Most genes exhibited some dependence on their mRNA (Fig. 1*D*,*E*). To incorporate transcript information, we assessed various low-complexity models involving one (linear), two (hyperbolic) or three (3-parameter logistic or HL) free parameters. The four models were compared by BIC, and HL was overwhelmingly the best or near-best model for most genes (Fig. 2*B*). Results were similar when using BIC weights (34) to assess the relative likelihood of HL against the others (*SI Appendix,* Fig. S2*D*). HL accommodated rare log-concave down relationships that occurred when protein abundance saturated at high transcript abundances (*p* < 10^-15^; Fig. 2*C*). Loss of transcript dependence arises biologically when a protein subunit surpasses the abundance of the complex in which it resides (21, 35), as observed for M genes across their entire measured range of mRNA. Accordingly, among HL genes, those that were log-concave down were mildly enriched in protein complexes in the CORUM database (*p* < 0.05) (33). More common were log-concave up HL relationships in which protein abundance increased only at higher mRNA abundances (Fig. 2*D*). Using a simple computational model for synthesis and turnover of mRNA and protein with dimensionless rate-parameter estimates, we found that log-concave up relationships arose naturally when steady-state abundances of mRNA and protein were halved and randomly sampled along the trajectory back to steady state (*SI Appendix,* Fig. S2*E,F*). Such “halving-and-random-sampling” occurs when cells asynchronously undergo cytokinesis, halving the protein copies per cell and sporadically re-entering into G1. Other HL genes showed mixed concavities or were linear to different extents (*SI Appendix,* Fig. S2*G–I*). Taken together, HL regressions provided the flexibility needed to capture various biological mechanisms that relate mRNA to protein (Fig. 1*D*).

One-third of HL regressions were statistically improved by adding mRNA features selected and weighted using LASSO (Fig. 1*E*). LASSO features were enriched for cytoplasmic translation (GO:0002181; *q* < 10^-10^) and the proteasomal pathway (GO:0043161; *q* < 10^-10^), suggesting dependencies that may promote protein synthesis or turnover. Specific *RPS*- or *RPL*-prefixed transcripts of the ribosome and *PSM*-prefixed transcripts of the proteasome were also enriched among LASSO-selected features (*p_ribosome_* < 10^-109^, *p_proteasome_* < 10^-18^). Ribosome feature weights were disproportionately positive, whereas proteasome subunits were negative (Fig. 2*E*), consistent with their expected influence on protein abundance. Overall, we asked whether LASSO-selected genes were more likely to interact physically with the protein of interest. Using the STRING database (36), we found a remarkable enrichment for interactions among LASSO-selected genes (*p* << 10^-4^; Fig. 2*F*), indicating that features contain more than spurious statistical associations. We conclude that Pinferna’s three-tiered modeling approach captures various biological phenomena and mechanisms in its framework.

### Pinferna predictions in cell lines and tissues relative to competing alternatives

To be useful for new samples, gene-specific model predictions should be more accurate than guesses based on past copy-number estimates of the protein in other settings. Therefore, we assessed Pinferna predictions against a null model built by iteratively drawing randomized measurements for each protein’s abundance from the meta-assembled dataset originally used for training (Fig. 1*A*). Accuracy was quantified by subtracting the measured value from the predicted value for each protein, taking the absolute value, and dividing by the standard deviation of the individual protein abundance across the 369 cell lines in the training data. This variance-scaled residual inversely weighs error by the breadth of abundances observed in other biological contexts. Finally, we compared the distribution of variance-scaled residuals between Pinferna and 100 null models to arrive at a proteome-wide estimate of model performance.

The first accuracy test was performed with HeLa cells, a line excluded from one of the original meta-assembled resources. We leveraged an independent study that carefully examined HeLa-to-HeLa differences with paired transcriptomics and SWATH proteomics (37). Pinferna consistently outperformed randomized measurements for all 12 HeLa derivatives investigated (*p* < 10^-6^; Fig. 3*A* and *SI Appendix,* Fig. S3*A*). Accuracy estimates were comparable when reported protein abundances were used instead of protein re-quantifications performed exactly as done for the training data (*Materials and Methods*; *SI Appendix,* Fig. S3*A,B*). Pinferna predictions were similarly resilient to reductions in transcriptomic sequencing depth—accuracies were comparable down to about 500,000 reads and remained superior to randomized measurements until about 50,000 reads (*SI Appendix,* Fig. S3*C*). The results bolster recent claims that typical single-cell RNA-seq data sequenced at ∼50,000 reads per cell poorly reflect protein abundances (38, 39) and separately indicate that Pinferna’s bulk predictions of protein from mRNA are robust to algorithmic details.

**Fig. 3.**
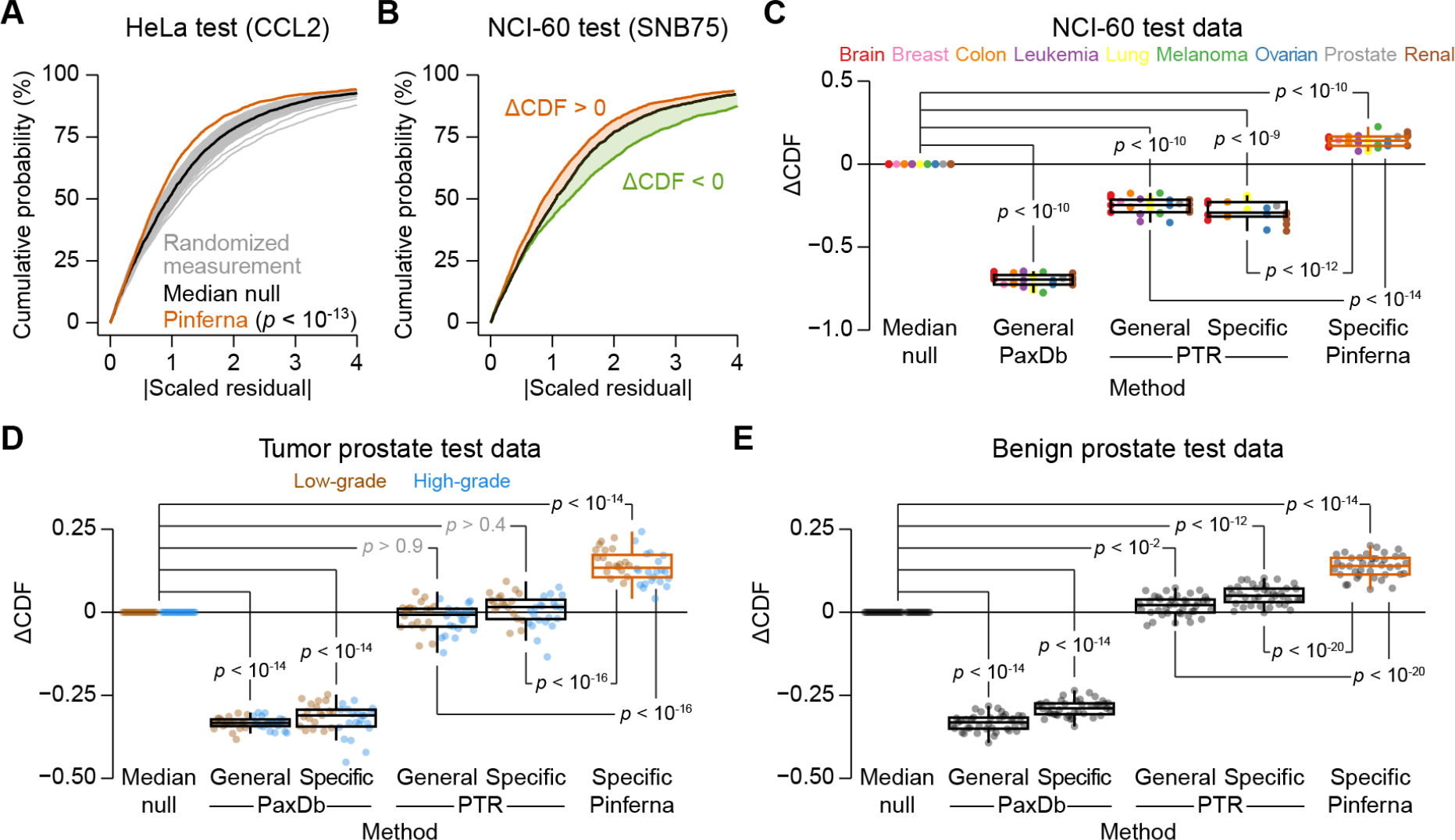
Pinferna outperforms empirical guesses and competing methods for absolute protein abundance estimation. (*A*) Pinferna compared to random protein-specific guesses. Model predictions were nondimensionalized as a scaled residual by subtracting the measured abundance, dividing by the standard deviation of the SWATH-scaled protein measured across the meta-assembly, and taking the absolute value (|Scaled residual|). The |Scaled residual| cumulative density was compared to randomized measurements drawn from the SWATH-scaled proteomic data for each gene. Randomized measurements were iterated 100 times (gray) to identify a median null (black) that served as a null distribution for model assessment. Left-shifted distributions indicate improved proteome-wide accuracy (relative to each protein’s variability) compared to protein-specific randomized measurements. Pinferna predictions of HeLa cells (orange; PRJNA437150; PXD009273) were compared to the null distribution by K-S test (*p* < 10^-13^). (*B*) Aggregate performance assessment of protein abundance predictions. The difference in cumulative density functions between test predictions and the median null distribution (ΔCDF) was integrated to identify approaches that performed better (ΔCDF > 0, orange) or worse (ΔCDF < 0, green) than protein-specific guessing. Data are from a prediction of Pinferna (orange) and tissue-specific protein-to-mRNA ratio (PTR; green). (*C–E*) Pinferna is consistently and uniquely superior to empirical guessing. ΔCDF values were calculated for NCI-60 cell lines (*C*; PRJNA433861; (43)) excluded from model training (Fig. 1A) and organized by cancer type (*n* = 5 brain, 1 breast, 3 colon, 4 leukemia, 4 lung, 3 melanoma, 3 ovarian, 1 prostate, 5 renal), primary prostate cancer samples organized by grade of the cancer (*D*; *n* = 19 low-grade, 21 high-grade), and normal prostate tissue (*E*; *n* = 39; PRJNA579899; PXD004589). PaxDb (40) and PTR (41) were used generically or in a tissue-specific way as alternative approaches (*Materials and Methods*). A cell line-specific PaxDb estimate was only available for U251 cells. Differences between groups were assessed by rank-sum test with Šidák correction. Box-and-whisker plots show the median (horizontal line), interquartile range (IQR, box), and an additional 1.5 IQR extension (whiskers) of the data.

To test Pinferna more broadly against other methods for protein estimation, we integrated the difference between the cumulative distribution functions of a model prediction and the median null to yield a single measure of accuracy improvement (ΔCDF; Fig. 3*B*). By ΔCDF, Pinferna was compared with two alternative approaches: 1) PaxDb, a meta-repository of protein abundances largely determined by uncalibrated peptide- and spectral-counting methods (40); and 2) protein-transcript ratios (PTR), which proportionately relate mRNA to protein abundances from data collected in 29 tissues (41). PaxDb and PTR each accommodate generic predictions using all available data and specific predictions restricted to data from a cell line or tissue of interest; both implementations were tested when possible. For the comparative evaluation, we assembled transcriptomic and SWATH data from 29 cell lines of nine cancers from the NCI-60 panel not included in the meta-assembly (42, 43). Overall, Pinferna was significantly more accurate than randomized measurements, whereas PaxDb and PTR were less accurate (Fig. 3*C*). Results were unchanged when transcriptomics were pre-processed with a different alignment pipeline (*SI Appendix,* Fig. S3*D*), reinforcing that Pinferna is tolerant of how mRNA TPMs are calculated. We observed no bias in estimates among cancer types (Fig. 3*C*) and thus concluded that Pinferna was the preferred method for predicting protein copy numbers from mRNA in cultured cell lines.

The performance in cell lines prompted us to ask whether Pinferna estimates would hold in more complex samples such as tissue. We assembled transcriptomic and SWATH data for 40 primary tumors and 39 normal human prostate samples (44). PaxDb remained significantly worse than randomized measurements (*p* < 10^-14^; Fig. 3*D,E*), consistent with the recognized limitations of peptide and spectral counts (11, 45). PTR was on par with randomized measurements in prostate cancer samples and significantly better in normal prostate samples, likely due to its tissue-centric focus (41). Nevertheless, Pinferna remained well ahead of all methods, indicating generality of proteome-wide copy number estimates from transcriptomics in primary tumors and nonmalignant tissues.

### Application to in silico modeling

RNA-seq often substitutes for protein when parameterizing systems-biology models of signaling (9), metabolism (8), and cell fate (10). For example, in constructing a mass-action model (24) of cardiomyocyte infection by coxsackievirus B3 (CVB3), RNA-seq was used to estimate abundances of the serial CVB3 receptors, CD55 and CXADR (Fig. 4*A*). Both estimates were HL extrapolations from a very-limited set of SWATH–RNA-seq pairings, which motivated a direct assessment of protein abundance by quantitative immunoblotting with recombinant standards (24, 46). Direct protein estimation was feasible for cultured cell lines but would be impossible for human hearts, where the severity of CVB3 infections is highly variable (47). We reasoned that Pinferna could provide extensibility to address this challenge and similar needs in cancer (8–10) and neurologic disease (20).

**Fig. 4.**
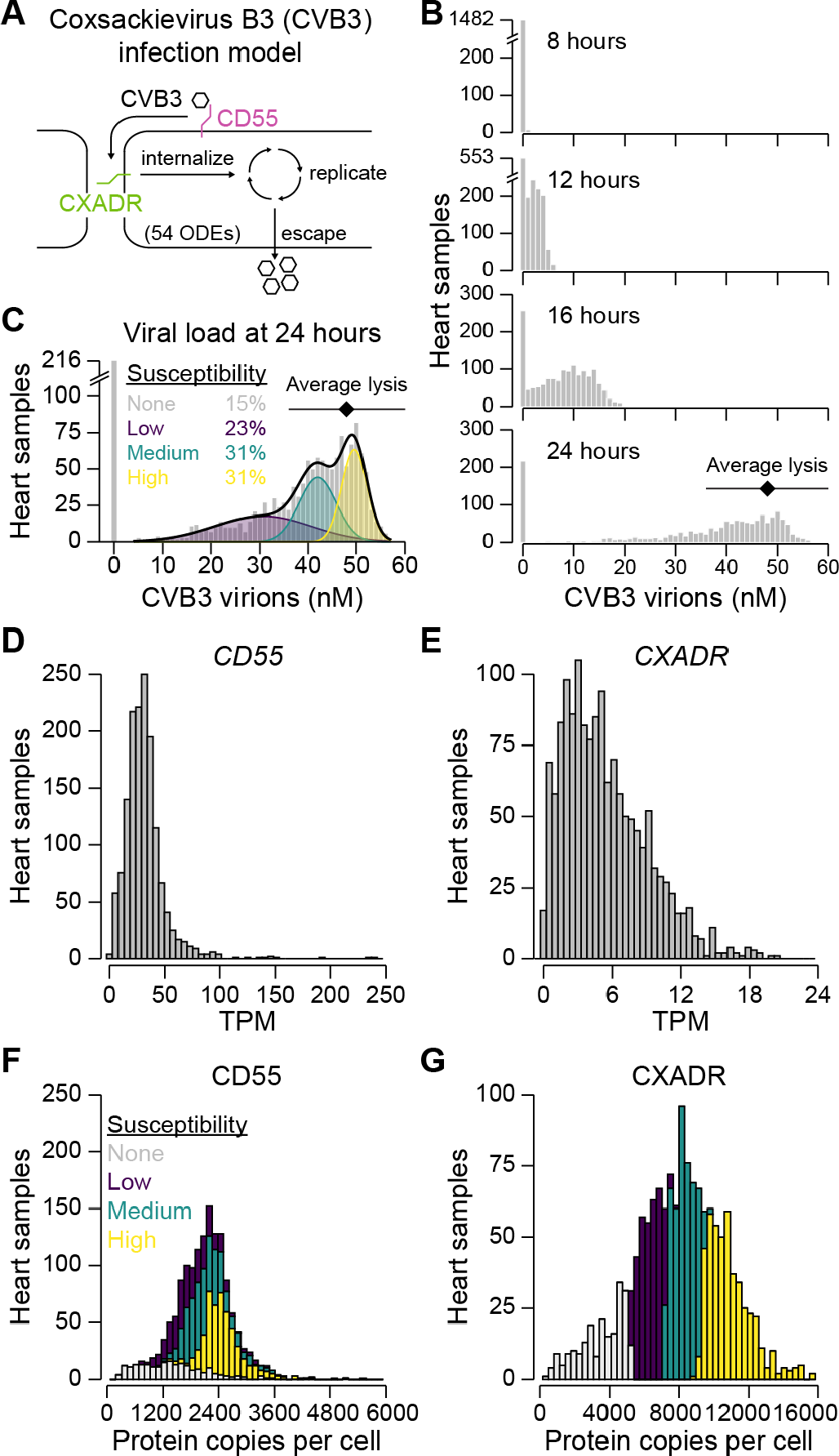
Simulating degrees of human cardio-susceptibility to coxsackievirus B3 (CVB3) infection based on inferred abundance differences in CVB3 receptors. (*A*) An in silico model of CVB3 initiated by its receptors CD55 and CXADR. After binding, the virus undergoes internalization, replication, and escape. The viral life cycle is mathematically modeled with 54 ordinary differential equations (ODEs; MODEL2110250001). (*B*) Distribution of viral load over time from 1489 human heart samples. Inferred abundances of CD55 and CXADR from each sample were used to simulate CVB3 infection. Each model run consisted of 100 simulated infections up to 24 hours with a coefficient of variation in model parameters of 5%. Viral loads (gray) at the indicated time points are shown along with the estimated point of lysis (black: mean estimated lytic yield ± s.d. (24, 25, 53)). (*C*) Four modes of infection susceptibility to terminal CVB3 infection. Viral load at 24 hours was replotted from *B* fit to a Gaussian mixture model (black) of three components (purple, green, yellow). Relative population densities in each of the susceptibility groups is shown along with the estimated point of lysis (black: mean estimated lytic yield ± s.d. (24, 25, 53)). (*D, E*) Distribution of mRNA abundances for *CD55* (*D*) and *CXADR* (*E*) normalized as TPM. (*F, G*) Distribution of inferred protein copy numbers per cell for CD55 (*F*) and CXADR (*G*) with each sample colored by its susceptibility.

RNA-seq data was collected for 1489 healthy and failing human heart samples from the U.S. and Europe, available through GTEx (48), MAGNet (49), and EGA (50). RNA-seq reads from the three studies were realigned and assembled together to enable comparability (*Materials and Methods* and *SI Appendix*, Table S8). The realignment confirmed no biases in expression based on data source for *CD55* (*SI Appendix*, Fig. S4*A*). *CXADR* expression increases in cases of cardiomyopathy (51, 52), and we reproduced this result by stratifying cases from the three sources (*p* < 10^-244^; *SI Appendix*, Fig. S4*B*). For both genes, the range of expression in heart samples fell within the variation observed across cancer cell lines in the meta-assembly (*SI Appendix*, Fig. S4*C,D*). CD55 is an HL+LASSO gene whose predictions are conditionally dependent on nine other genes. These features reduce CD55 inferences below the smoothed average of the training at very low TPM (*SI Appendix*, Fig. S4*C*). By contrast, CXADR is an HL gene that is steeply nonlinear at low TPMs where small changes greatly influence the protein copy number estimate (*SI Appendix*, Fig. S4*D*). Because of the nonlinear inferences, CD55–CXADR proteins were strongly coupled even while *CD55*–*CXADR* mRNAs were much less so (*p* < 10^-12^; *SI Appendix*, Fig. S4*E,F*). Using the Pinferna estimates of CD55 and CXADR as initial conditions, we created a series of individualized model variants to create a virtual cohort of the human population. The individualized models were initiated with a high titer of CVB3 that guaranteed infection of permissive cells, and the concentration of virions was collected at each time point for 24 hours of simulated infection (*Materials and Methods*). The goal was to investigate whether inferred CD55–CXADR protein variations yielded a wide enough range of infection outcomes in the model that one or both receptors could be nominated as a susceptibility factor.

Examining the predicted distribution of viral loads over time, we noted a strong asymmetry in onset of infection (Fig. 4*B*). At 12 hours, 64% of individuals were detectably infected, producing mature virions above one plaque forming unit (1 pfu = 0.48 ± 0.12 nM in these simulations). By 24 hours, the models yielded a left-skewed distribution, which straddled the mean lytic yield of viruses in the CVB3 genus (∼100 pfu = 48 ± 12 nM for a 3700 µm^3^ cell) (24, 25, 53). Even at this uncharacteristically late time, 15% of individuals remained uninfected, suggesting they were intrinsically resistant. The remaining cases were best fit as a three-component Gaussian mixture of low, medium, and high susceptibilities (Fig. 4*C*). Based on mean lytic yield, we interpreted these groups as prone to subinfection, infection, and severe infection, with failing hearts falling almost entirely into the infection and severe-infection groups (*SI Appendix*, Fig. S4*G*). For comparison, we abandoned Pinferna and attempted a randomized-measurement approach by linearly scaling the RNA-seq data about a quantity of CD55 and CXADR arbitrarily selected from the training data (*Materials and Methods*). As expected, model outputs were so dependent on the randomized measurement that they were uninterpretable when viewed in aggregate (*SI Appendix,* Fig. S4*H*). Randomized measurements tended to predict ∼100% resistance or ∼100% lytic infections and underestimate the low-susceptibility group, although some fortuitously matched the true inferences. We concluded that the Pinferna-derived model outputs were compelling enough to interpret further.

Among heart samples, the distributions of *CD55* and *CXADR* RNA transcripts were quite different (Fig. 4*D,E*). The range of *CD55* expression was ∼tenfold that of *CXADR*, hinting that it might be the dominant receptor for in silico susceptibility. However, these population-wide trends changed when viewed as protein inferences (Fig. 4*F,G*). Both CD55 and CXADR were more symmetrically distributed, with CXADR exhibiting greater overall variance. Importantly, when individuals were classified on the basis of their inferred susceptibility, we found that CXADR abundance alone was sufficient to stratify the population. This application of Pinferna illustrates how direct substitution of transcriptomics can misconstrue the outputs of systems-biology models built for protein networks.

### Application to molecular subtyping

Transcriptomic profiles are widely used to define disease subtypes (54–56), which may change when gene expression is replaced by inferred protein abundance as a closer surrogate of cell function. As a longstanding example, we selected the intrinsic molecular subtypes of breast cancer defined by a 50-gene classifier (PAM50) for 796 cases with RNA-seq in The Cancer Genome Atlas (26, 57, 58). For consistency, our analysis focused on the 4366 transcripts compatible with protein inference (Fig. 1*A*), but results were unchanged when using the entire available transcriptome (*SI Appendix,* Fig. S5*A–D*). Consensus clustering of mRNA profiles identified five ordered and stable groups, which were statistically enriched in PAM50-assigned cases of 1) Normal-like, 2) HER2+, 3) Luminal A, 4) Basal-like, and 5) Luminal B breast cancer (*p* < 0.004 by hypergeometric test; Fig. 5*A* and *SI Appendix*, Fig. S5*A,B*). When the analysis was repeated with Pinferna estimates after standardization, the smallest number of stable and significant consensus clusters was again five (*SI Appendix*, Fig. S5*E,F*). However, the enriched PAM50 assignments were reordered, and 186/796 = 23% of cases changed to a different cluster (Fig. 5*A*). The aggregate transformations of Pinferna (Fig. 1*C–E*) thus exceeded a standardized rescaling and considerably altered subgroup composition.

**Fig. 5.**
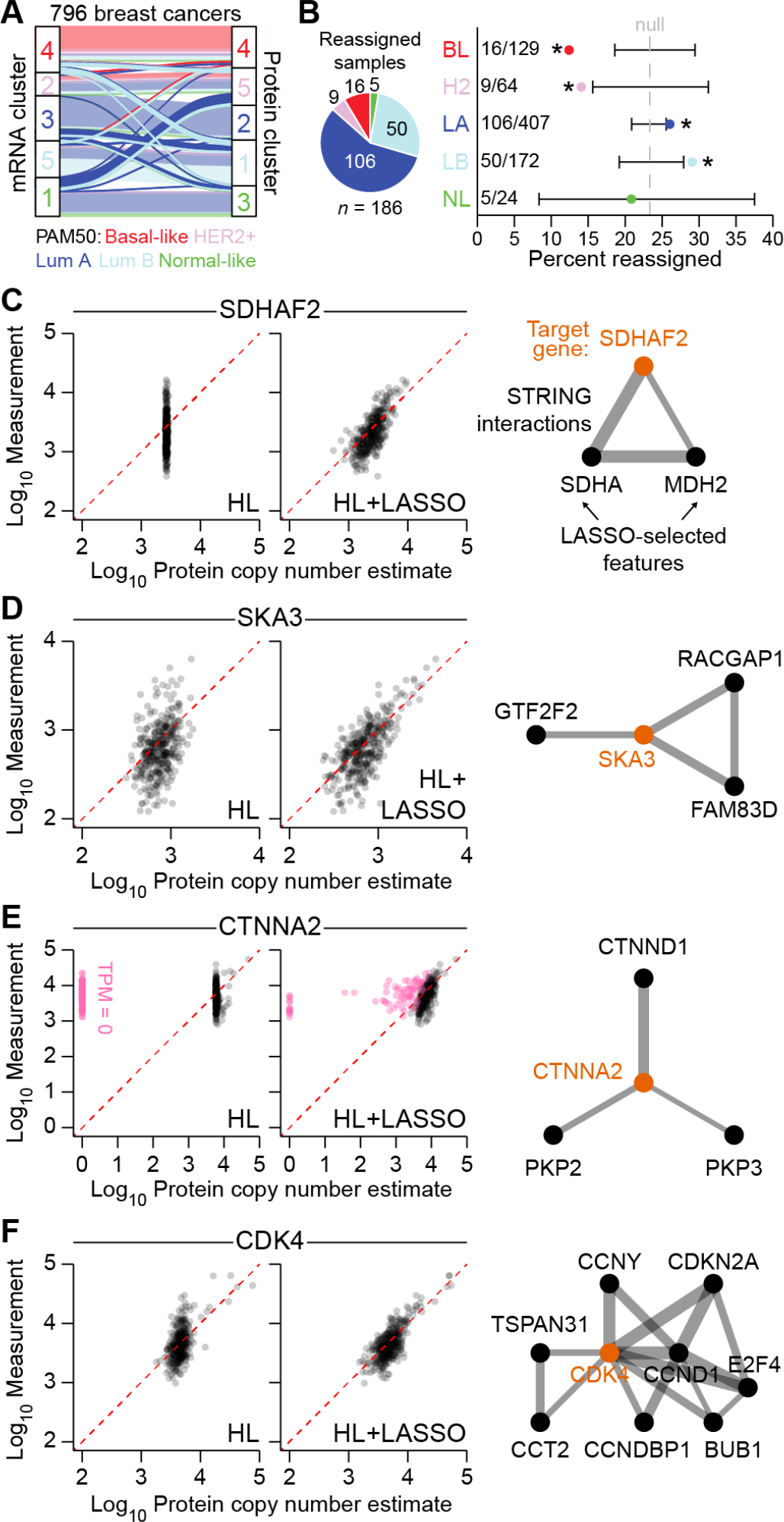
Inferred proteomics reassigns luminal A/B transcriptomic subtypes of breast cancer. (*A*) Reorganization of five consensus clusters defined by RNA-seq (left) and Pinferna (right) for 796 breast cancers in The Cancer Genome Atlas (26). Clusters were determined by Monte Carlo consensus clustering (91) and colored according to the dominant PAM50 subtype of each cluster. Samples that did not change clusters are transparent in the background while samples that changed are opaque in the foreground. Lum A: Luminal A; Lum B: Luminal B. (*B*) Reassigned samples are predominated by luminal A/B PAM50 subtypes. (*Left*) Proportion of each subtype among samples that were reassigned. (*Right*) Percent reassignments for each subtype. The average overall reassignment rate is shown as a null reference (186/796 = 23%; gray dashed) with the 90% hypergeometric confidence interval (black) for each subtype. Reassignment enrichments were determined by hypergeometric test, asterisk indicates *p* < 0.05. BL: Basal-like; H2: HER2+; LA: Luminal A; LB: Luminal B; NL: Normal-like. *(C–F)* Cluster-reorganizing genes are highly dependent on other genes. (*Left*) Concordance between SWATH measurements and the HL fit ± LASSO in the meta-assembly. Perfect concordance is given by the red dashed line. Pink points in *E* are samples with TPM = 0 for *CTNNA2*. (*Right*) STRING interactions (edges) among the target gene (orange) and its LASSO-selected features (black). Edge thickness (gray) reflects the confidence of the interaction as determined by STRING. Thicker lines represent a higher confidence score. Line lengths are arbitrary.

Among reassigned samples, we noted preferential enrichments in Luminal A (26%) and Luminal B (29%) tumors over Basal-like and HER2+ (Fig. 5*B*). Luminal A/B cases are often intermingled in clusters defined by transcriptomics (59), prompting us to look more deeply at their reassignment with Pinferna (*SI Appendix*, Fig. S5*G*,*H*). We surveyed for HL+LASSO genes whose z-score standardized values changed the most from mRNA to inferred protein and looked within these influential genes for features (other genes) that were STRING interactors (Fig. 2*F* and *SI Appendix*, Fig. S5*I*; *Materials and Methods*). Calibration of the mRNA-to-protein relationship for SDHAF2 (a mitochondrial Complex II assembly factor) was dramatically improved with abundance information from other genes, including its interactors, SDHA and MDH2 (Fig. 5*C*). Similarly, inference of SKA3 (a subunit of the mitotic Ska complex) was influenced by multiple binding partners (Fig. 5*D*). One of the most notable examples of LASSO modulation was CTNNA2 (an adhesion protein involved in actin regulation). CTNNA2 protein was ubiquitous in the meta-assembly, but its mRNA was undetectable in 23% of training samples; of these, nonzero protein inference of 83% was recovered by using abundances including CTNND1 and plakophilin (PKP2, PKP3) interactors (Fig. 5*E*). For cluster reassignments of breast cancer, the gene with the most interaction-rich feature set was CDK4. Like SDHAF2, CDK4 protein abundance was largely independent of its own mRNA, but a useful calibration was achieved when considering various cyclins and other binding proteins (Fig. 5*F*). This result is important because CDK4/6 inhibitors are approved to treat luminal breast cancers (60, 61), but responsiveness has not consistently associated with the abundance of *CDK4* mRNA (62, 63). Newly meaningful subtypes with therapeutic implications might arise when combining transcriptomics with Pinferna to get closer to the functional proteome.

## Discussion

We have devised a straightforward gene-by-gene formalism that uses mRNA to achieve absolute protein estimates informed by the best measurements available for each data type. Gene-specific inferences are gleaned from cancer cell lines, but general accuracy and utility is verified in multiple other contexts. Although not exhaustive, our coverage of 4366 mRNA–protein relationships is considerable given that modern SWATH experiments reliably quantify ∼5000 proteins (37). Encouraged by the robustness of predictions to read depth and alignment details, we provide Pinferna as an open resource (http://janeslab.shinyapps.io/Pinferna; *SI Appendix*, Table S9). The platform is optimal when provided full RNA-seq profiles, but it also accommodates single-gene TPM entries for LASSO-free inference when subset data are exported from public repositories. The hope is to attract general users to the value in seeing genes and gene profiles of interest through the lens of inferred proteomics (source code is available for developers; *Materials and Methods*).

The mRNA-to-protein relationship classes used here distill the major findings of prior models that were more fine grained (41, 64, 65). Measured mRNA is the net result of its transcription– degradation, with the per-mRNA yield of translation for each gene being the greatest determinant of absolute protein abundance (17, 65). Together, mRNA abundance and per-mRNA yield define an expected set point for protein abundance as captured by the HL relationship class. For typical in vitro cultures, cell doubling is faster than turnover of most proteins (19, 65), creating a perpetual state of halving-and-recovery that likely explains why many HL models are log-concave up. Nonetheless, overall accuracy of Pinferna did not decrease with clinical samples that were less proliferative, suggesting a role for other cyclic perturbations in vivo, such as circadian rhythms (66). Some genes additionally require protein complexes to persist stably (19, 21), which creates buffering dependencies on other genes that are coexpressed. The HL+LASSO approach seeks to capture this relationship class by identifying statistical mRNA–protein associations in trans. Despite its heavy L1 regularization (67), LASSO recovered a significant number of documented protein-protein interactions. Recently, a reciprocal approach to predict relative protein abundance from mRNA was proposed that constrains the search space for each gene to its CORUM–STRING interactors but relaxes the regularization by using elastic net (30, 67). These models retain many more features (158–457 (30) compared to 1–83 features for HL+LASSO), but their relative predictions cannot be compared with the absolute copy-number estimates of Pinferna. Future versions may consider hybrid regularizations that penalize CORUM– STRING interactors less during LASSO feature selection. Iterative approaches might also use Pinferna inferences as LASSO features for other genes to approximate biological dependencies more closely. Lastly, we speculate that M genes arise from protein complexes so large that pairwise interactions within a complex completely dictate abundance (68, 69).

Absolute protein estimates are of great practical use for making qualitative determinations in systems-biology models. For example, using inferred protein to simulate heart infections, we clarified that individuals with fewer than ∼5000 copies of CXADR per cell were not susceptible to CVB3 (Fig. 4*G*). Another subfield of relevance is genome-scale metabolic modeling with tailored derivatives of the generic human metabolic network reconstruction (70). For cell- or tissue-specific modeling, the generic reconstruction is pruned according to which metabolic genes are “not expressed” in a biological context of interest (71, 72). Irrespective of the pruning algorithm, the choice of threshold is made absolutely across all genes in a sample, which defines the resulting model complexity (73). Protein abundances for some metabolic pathways scale linearly with mRNA, but others do not (6, 7), and protein–mRNA set points vary over several orders of magnitude (17, 41). For metabolic models concerned with protein fidelity, estimating copy numbers from RNA-seq is a scalable alternative to proteome immunohistochemistry (72).

There are limitations in our approach to absolute copy-number estimation. By relying on SWATH for calibration, we lose many of the 8000+ proteins quantified in relative terms by TMT (27). Calibration data were all collected at steady state; thus, we caution against using RNA-seq obtained shortly after acute perturbations when transcripts and proteins will be most uncoupled (19). Signaling proteins rapidly turned over by ubiquitylation (TP53, NFE2L2, NFKBIA) might require other formalisms when they become detectable by SWATH. Broadening Pinferna predictions to non-human samples awaits the availability of robust SWATH libraries in other mammals (74). Last, we recall that no total-protein estimator captures functional state, such as the surface localization of CD40, the tyrosine phosphorylation of PKP2–PKP3, or the kinase activity of CDK4 (Fig. 2*C*, 5*E*,*F*). Despite these caveats, the method adds an immediately useful approach for systems biology that compares favorably against existing alternatives.

In statistics, bootstrapping by computation allowed the field to tackle analytical needs that were impossible to address by existing methods (75). Analogously, proteomics can extend its reach by combining with RNA-seq to inform more biological samples, including retrospective ones it would never otherwise have access to. There is already precedent for such “transcriptomic bootstrapping” to infer genomic alterations (76), molecular kinetics (76), and cellular trajectories (77).

## Materials and Methods

### SWATH alignment and quantification

Raw SWATH data files were obtained from the PRIDE repository (15) (CAL51, PXD003278 (28); U2OS, PXD000954 (29); HeLa, PXD00927 (37)) and converted to .mzML format using the MSConverGUI (version 3.0) in the ProteoWizard software suite (78) with the following options: Output format, mzML; Extension, mzML; Binary encoding precision, 64-bit; Write index; Use zlib compression. Peptide fragments were aligned with OpenSwathWorkflow in OpenMS (version 2.4.0) (79) with the following options: -sort_swath_maps, -readOptions normal, - batchSize 1000, -use_ms1_traces, -mz_correction_function quadratic_regression_delta_ppm. Statistical control was performed with PyProphet (version 2.2.5) (80) with the following options: -- group_id=transition_group_id, --tric_chromprob. Each series of SWATH runs was realigned with TRIC (81) using msproteomicstools (version 0.11.0) with the following options: --method LocalMST, -- realign_method lowess, --max_rt_diff 60, --mst:useRTCorrection True, --mst:Stdev_multiplier 3.0, -- target_fdr 0.01, --max_fdr_quality 0.05, --alignment_score 0.0005. The top three peptide fragments by intensity (or all peptide fragments if fewer than three) were summed for each protein to estimate relative abundance. Summed intensities were mean-averaged across technical replicates when available. To place summed intensities on an absolute scale, the median abundance of all detected proteins within each sample was centered at 10,000 protein copies per cell (37).

### RNA-seq alignment and quantification

For all studies other than Fig. 4, SRA files were obtained from the Sequence Read Archive (SRA) (82) (HeLa, PRJNA437150; NCI-60, PRJNA433861; Prostate, PRJNA579899) and converted to raw FASTQ files using sratoolkit (version 2.10.5) with fasterq_dump. TruSeq adapters were trimmed using the fastq-mcf function in the ea-utils package with the following options: -q 10, -t 0.01, -k 0. Trimmed datasets were aligned to the human genome (GRCh38) using HISAT2 (version 2.1.0) (83) with the following options: --dta (downstream transcriptome assembly) and either --rna-strandedness RF (for paired-end reads generated by the TruSeq strand-specific library; NCI-60 and prostate samples) or --rg-id (for single-end reads generated by the TruSeq library; HeLa). Output SAM files were converted to BAM files using the sort function in samtools (version 1.12) (84), and BAM files were indexed to create BAI files using the index function for obtaining counts downstream. Alignments were assembled into transcripts using StringTie (version 2.1.0) (85) with the - e option restricting assembly to known transcripts in the provided annotation. Counts were obtained using HTSeq (version 2.0.2) (86) using BAM files as the input with the following options: -f bam, -r pos, -m intersection-strict, -s reverse, -a 1, -t exon, -i gene_id. Heart RNA-seq data were obtained from dbGaP (GTEx, phs000424.v9.p2), the Sequence Read Archive (MAGNet, SRP237337), and the European Genome-Phenome Archive (EGA, EGAS00001002454). GTEx samples were converted from BAM files to raw FASTQ files using samtools (version 1.12) (84). MAGNet samples were converted to raw FASTQ files using sratoolkit (version 2.10.5) (https://hpc.nih.gov/apps/sratoolkit.html#doc). EGA samples were converted to raw FASTQ files using samtools (version 1.10/1.12) (84) and paired reads were recreated using the fastqCombinePairedEnd.py script from Eric Normandeau (https://github.com/enormandeau/). Datasets were aligned to the human genome (GRCh38) using HISAT2 (version 2.1.0) (83) with the --dta option for downstream transcriptome assembly. Output SAM files were converted to BAM files using samtools (version 1.12) (84). Alignments were assembled into transcripts using StringTie (version 2.1.0) (85) with the -e option restricting assembly to a unified list of transcripts that was provided by first running StringTie using the --merge option.

### Data harmonization

The table of MANE Select identifiers was obtained from the source publication (87) and filtered for “MANE Select” genes. The filtered table was appended with UniProt accession codes using biomaRt (version 2.52.0) and GRCh38. The Ensembl BioMart browser was used to obtain HGNC identifiers, Ensembl transcript identifiers (with version numbers for maximum overlap), RefSeq mRNA identifiers, NCBI (formerly Entrez) gene identifiers, UniProt accession codes, and UniProt gene symbols for *Homo sapiens*. Each row of the MANE Select table was matched to at least two identifiers in the biomaRt table to determine the UniProt accession numbers. When MANE Select annotated a gene symbol as LOC###### and biomaRt contained a more descriptive gene symbol, the biomaRt gene symbol replaced the MANE Select gene symbol and the “Database” column was updated to include “biomaRt symbol” as the source. The harmonization identified 83 genes that are not currently available in UniProt. The final harmonized table of ten identifiers (nine for the 83 genes not in UniProt) for 19,062 genes is available in *SI Appendix*, Table S2.

### Cancer Cell Line Encyclopedia pre-processing

The TMT proteomic dataset was obtained from the source publication (27) as a CSV file (protein_quant_current_normalized.csv). After removing proteins annotated as “Fragments”, gene symbols were matched to UniProt accession codes by using the harmonized identifier table (*SI Appendix*, Table S2). Protein isoforms with redundant gene symbols were summed. The RNA-seq dataset (31) was obtained from the Depmap portal, back-transformed from log_2_ to TPM, and renamed with the harmonized identifier table (*SI Appendix*, Table S3).

### Meta-assembly, calibration, and inference

#### 1. Scaling

For each gene, SWATH copy-number estimates were divided by the corresponding harmonized TMT data for U2OS and CAL51 cells to calculate U2OS- and CAL51-specific scaling factors. Scaling factors were averaged when possible; otherwise, a single scaling factor was used (*SI Appendix*, Fig. S1*A*). The resultant scaling factors were then multiplied across the harmonized TMT data table to yield a SWATH-scaled proteomics dataset of 4385 total proteins across 375 cell lines (*SI Appendix*, Table S1).

#### 2. Regression

The SWATH-scaled proteomics and RNA-seq transcriptomics datasets were filtered before regression. As recommended (88), proteomics data from replicates of CAL120 (CAL120_BREAST_TenPx02), SW948 (SW948_LARGE_INTESTINE_TenPx11), and HCT15 (HCT15_LARGE_INTESTINE_TenPx30) were excluded. RNA-seq data were filtered to include only cell lines with SWATH-scaled proteomics available. Both datasets were filtered to retain genes for which SWATH-scaled proteomics was available in at least 150 cell lines (4445/4513 = 98.5% of all SWATH-scaled proteins). Filtered datasets are available in *SI Appendix*, Tables S3 and S4. Numerical approaches for building the M, HL, and HL+LASSO models and assessment of confidence intervals is described in *SI Appendix*, *Materials and Methods*.

#### 3. Model selection

The BIC for each regression was calculated under the assumption of normally distributed random errors as follows:

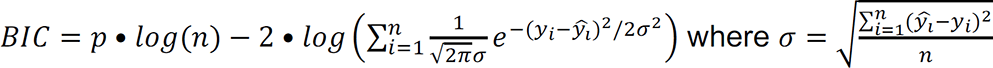

Where *σ* is the standard deviation of the fit, *n* is the number of observations, *p* is the number of model parameters, 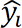 is the predicted value of the *i*^th^ observation, *y_i_* is the measured value of the *i*^th^observation, and *LL* is the log-likelihood with *log* being the natural logarithm. Comparison of HL with linear, hyperbolic, and 3-parameter logistic alternatives is described in *SI Appendix*, *Materials and Methods*.

### Gene ontology analysis

Enrichments of M, HL, and HL+LASSO genes for biological processes were evaluated with the GO knowledgebase (89).

### Concavity analysis

The concavity of HL fits was assessed with the check_curve function of the inflection package (version 1.3.6) in R. HL curves for TPM > 5 (∼1 copy per cell) were analyzed after log transformation of x and y coordinates.

### Feature weight distributions

To obtain LASSO feature weights, each LASSO coefficient for a gene was multiplied by the mean TPM of the feature averaged across the 369 cell lines in the meta-assembly (including zeros). Feature weights for all HL+LASSO genes were concatenated and filtered for subunits of the proteasome (*PSM*-prefixed gene names) or ribosome (*RPS*- or *RPL*-prefixed gene names), allowing duplicates if the feature appeared in more than one gene. Distributions were plotted as smoothed densities with the geom_density function in ggplot2 (version 3.4.0).

### STRING interactions and maps

STRING interactions were obtained with the STRINGdb (version 2.8.4) and rbioapi (version 0.7.7) packages in R. Sessions were initialized with STRINGdb$new and the following arguments: species = 9606 (Homo sapiens), version = 11.5, score_threshold = 400 (medium-confidence interactions). HL+LASSO genes were mapped with the string_db$map function, and up to 1000 medium-confidence interactions were retrieved with the rba_string_interaction_partners function and harmonized with the gene identifier table (*SI Appendix*, Table S2). For comparison, LASSO-selected features were substituted with an identical number of randomly selected genes to recalculate the number of interactions. The substitution–recalculation step was iterated 10^4^ times to build a null distribution. Details about STRING visualizations are described in *SI Appendix*, *Materials and Methods*.

### Assembly of test datasets

#### HeLa

Using the latest analytical procedures, SWATH data from two HeLa derivatives [Kyoto L8 and CCL2 L13] in the original study (37) did not pass the internal calibration step of the OpenSwathWorkflow alignment and were omitted here. Pre-quantified SWATH data for HeLa derivatives were downloaded from https://helaprot.shinyapps.io/crosslab/ and normalized to the median copy number of proteins co-quantified in the meta-assembly (10,000 copies per cell). Downsampling of HeLa RNA-seq reads is described in *SI Appendix*, *Materials and Methods*.

#### NCI-60

Pre-aligned SWATH data for the NCI-60 panel of cell lines (43) were downloaded from CellMiner as a processed data set (Protein: SWATH (Mass spectrometry) - Peptide), quantified for protein, harmonized as described above, and normalized to the median copy number of proteins co-quantified in the meta-assembly (14,000 copies per cell). Pre-aligned RNA-seq data for the NCI-60 panel of cell lines (42) were downloaded from CellMiner as a processed data set (RNA: RNA-seq - composite expression), summarized as TPM, and harmonized as described above. The following NCI-60 lines excluded from the meta-assembly were used as test data: SF268 (brain), SF539 (brain), SNB-19 (brain), SNB-75 (brain), U251 (brain), Hs 578T (breast), COLO 205 (colon), HCC2998 (colon), KM12 (colon), CCRF-CEM (leukemia), HL-60(TB) (leukemia), MOLT-4 (leukemia), SR (leukemia), EKVX (non-small cell lung), HOP-62 (non-small cell lung), HOP-92 (non-small cell lung), NCI-H322M (non-small cell lung), M14 (melanoma), Malme-3M (melanoma), MDA-MB-435 (melanoma), NCI-ADR-RES (ovarian), OV-CAR5 (ovarian), SK-OV-3 (ovarian), DU145 (prostate), ACHN (renal), RXF 393 (renal), SN12C (renal), TK-10 (renal), and UO-31 (renal).

#### Prostate

Pre-aligned SWATH data for normal and malignant prostate (44) were downloaded from PRIDE (PXD004589), quantified for protein, harmonized for gene names as described above, and normalized to the median copy number of proteins co-quantified in the meta-assembly (14,000 copies per cell). Harmonization of samples was more challenging because of different patient-coding schemes for the SWATH and RNA-seq datasets. We obtained metadata annotation for RNA-seq from the SRA Run Selector (PRJNA579899) and then reconciled these identifiers with the PXD004589 identifiers using a key personally communicated by Wenguang Shao (90). Tumor-normal pairs were retained in the harmonized dataset if the tumor grade annotations were consistent between SWATH and RNA-seq. The final patient annotations and cross-referencing key is available in *SI Appendix*, Table S7.

#### Heart

The GTEx test dataset from the v8 final data release consisted of 432 left ventricle and 429 atrial appendage autopsy samples from 561 healthy donors (48). Both GTEx heart tissue sites were considered separately in the analysis. The MAGNet test dataset consisted of 200 cardiomyopathy and 166 healthy control samples (49). The EGA test dataset consisted of 149 cardiomyopathy and 113 healthy control samples (50). After RNA-seq alignment as described above, the 1489 samples were concatenated without batch correction before the analysis.

#### Breast cancer

Pre-aligned RNA-seq data for ductal and lobular neoplasms in The Cancer Genome Atlas (TCGA) were downloaded from the Genomic Data Commons portal. The samples were intersected by TCGA identifiers with the published samples profiled by RNA-seq and classified by PAM50 (26), yielding 796 samples in the test dataset.

### Alternative methods for protein abundance estimation

#### PaxDb

The following aggregated proteomics data were downloaded from PaxDb (40) as averaged protein parts per million (ppm): for NCI-60 comparisons, H.sapiens - Cell line (Integrated); for general tissue comparisons, H.sapiens - Whole organism (Integrated); for prostate tissue comparisons, H.sapiens - Prostate gland (Integrated). PaxDb entries less than 0.01 ppm were excluded, and the filtered data were harmonized as described above. Last, each aggregated dataset was normalized to the median copy number of proteins co-quantified in the meta-assembly: for Cell line (Integrated), 8000 copies per cell; for Whole organism (Integrated), 8000 copies per cell; for Prostate gland (Integrated), 9000 copies per cell. Note that PaxDb does not use information from RNA-seq and thus makes a single prediction of protein abundance for each integrated context. Cell line-specific information was not available for NCI-60 lines other than U251.

#### PTR

Protein-to-mRNA ratios in the original publication (41) were not calculated on the scale of copies per cell. To convert, the protein abundances used for PTR estimation were normalized to the median copy number of proteins co-quantified in the meta-assembly (9000 copies per cell). Using the renormalized protein abundances, we rederived PTRs from the associated RNA-seq fragments per kilobase per million mapped reads (FPKM) as follows: PTR = log_10_(protein abundance) - log_10_(FPKM) (41). PTRs were calculated for each of 29 tissues and in a general manner by using the median PTR. NCI-60 predictions used the PTR specific to each cell line’s tissue of origin, which was not available for leukemic, breast, or melanoma lines; therefore, these lines were omitted from PTR predictions.

### Randomized measurements, median null model, and ΔCDF

Randomized SWATH measurement profiles were constructed by randomly sampling a gene-specific copy number estimate for each gene in the proteome and iterating 100 times without replacement. The 100 randomized-measurement distributions were compared to Pinferna and ordered by the K-S statistic, with the median distribution selected as the null model for formal K-S hypothesis testing. To compare prediction methods to the median null, the area between the two was integrated by difference (ΔCDF) using the AUC function in the DescTools (version 0.99.47) package in R.

### Monte Carlo consensus clustering (M3C)

Consensus clustering of the breast cancer dataset was performed with all genes (19,062) or all genes inferrable by Pinferna (4366). LASSO-modulated protein inferences below zero copies per cell were set to zero. All data were log_2_ transformed, and genes with zero variance were eliminated before Z-score standardization. Datasets were clustered using the M3C function from the M3C (version 1.18.0) package with the following options: removeplots = T, iters = 10, objective = “entropy”, clusteralg = “spectral”. Clustering statistics lie within the M3C object, and five consensus clusters were selected based on maximum or near-maximum cluster stability and significance. Other hierarchical clustering of the breast cancer dataset is described *SI Appendix*, *Materials and Methods*.

### Transcription-translation model

The system of ordinary differential equations was solved in MATLAB R2022a using ode15s. The simulation was performed 100 times allowing parameters to vary lognormally about their central nondimensionalized estimate with a coefficient of variation of 10%. After each simulation, 10 time points were chosen randomly and stored, for a total of 1000 points at the end of the simulation.

### CVB3 model

A mass-action model of CVB3 infection (MODEL2110250001) was modified from its published version (24) to accommodate CD55 and CXADR abundances as input parameters. Each simulated infection was initialized with abundances for CD55–CXADR and 10 plaque-forming units of CVB3. In silico infections proceeded for 24 hours and were iterated 100 times with 5% lognormal coefficient of variation between runs. The median virion output was stored, and overall viral load (measured as mature CVB3 virions and after excluding cases of zero viral load) was fit with a Gaussian mixture model using the Mclust function in the Mclust (version 6.0.0) package in R. The best mixture model by BIC was a three-component model of unequal variance, which classified each sample based on the probability of the sample falling into that component. For the randomized-measurement case (*SI Appendix*, Fig. S4*H*), protein estimates were obtained by setting the median TPM value of the heart samples to a randomly selected protein set point in the meta-assembly and linearly scaling the other samples around that point. This process was replicated 100 times to create a 1489 sample x 100 replicate matrix of randomized measurements for CD55 and CXADR. Each CD55–CXADR pair was passed to the CVB3 model and simulated with MATLAB (version R2022b) as before 100 times each for a total of 14,890,000 simulations, which were threaded to 100 cores over 10 nodes on the Rivanna high-performance computing cluster of the University of Virginia. Taking viral load at 24 hours of infection as the phenotype, we classified [CVB3 virions] = 0 nM as resistant, 0 nM < [CVB3 virions] < 36 nM as sublytic, and [CVB3 virions] ≥ 36 nM as lytic.

### Software availability

Code is available on GitHub for the Pinferna web browser (https://github.com/JanesLab/Pinferna/) and the transcription-translation model (https://github.com/JanesLab/TranscriptionTranslation/).

## Acknowledgments

We thank Jeff Saucerman for suggesting BIC weights, William Shao for testing the Pinferna web browser, the University of Virginia High-Performance Computing group for computational guidance, and Kenley Ellis for editing the manuscript. This study was supported by grants from the National Institutes of Health (U54-CA274499 to K.A.J.) and the David & Lucile Packard Foundation (2009-34710 to K.A.J.), a Cardiovascular Training Grant Fellowship (T32-HL007284 to A.J.S.), a Sture G. Olsson Fellowship (to A.J.S.), and a Human Frontier Science Program Fellowship (LT000469/2021-L to C.D.G.).

## References

1. R. Phillips, R. Milo, A feeling for the numbers in biology. Proc. Natl. Acad. Sci. U. S. A. 106, 21465–21471 (2009).

2. Z. Wang, M. Gerstein, M. Snyder, RNA-Seq: a revolutionary tool for transcriptomics. Nat. Rev. Genet. 10, 57–63 (2009).

3. T. Barrett et al., NCBI GEO: archive for functional genomics data sets--update. Nucleic Acids Res. 41, D991–995 (2013).

4. R. S. Fehrmann et al., Gene expression analysis identifies global gene dosage sensitivity in cancer. Nat. Genet. 47, 115–125 (2015).

5. Z. Duren, X. Chen, R. Jiang, Y. Wang, W. H. Wong, Modeling gene regulation from paired expression and chromatin accessibility data. Proc. Natl. Acad. Sci. U. S. A. 114, E4914–E4923 (2017).

6. B. Zhang et al., Proteogenomic characterization of human colon and rectal cancer. Nature 513, 382–387 (2014).

7. P. Mertins et al., Proteogenomics connects somatic mutations to signalling in breast cancer. Nature 534, 55–62 (2016).

8. J. E. Lewis, T. E. Forshaw, D. A. Boothman, C. M. Furdui, M. L. Kemp, Personalized Genome-Scale Metabolic Models Identify Targets of Redox Metabolism in Radiation-Resistant Tumors. Cell Syst 12, 68–81 e11 (2021).

9. E. J. Pereira et al., Sporadic activation of an oxidative stress-dependent NRF2-p53 signaling network in breast epithelial spheroids and premalignancies. Sci Signal 13, eaba4200 (2020).

10. A. Montagud et al., Patient-specific Boolean models of signalling networks guide personalised treatments. Elife 11, (2022).

11. N. Pappireddi, L. Martin, M. Wuhr, A Review on Quantitative Multiplexed Proteomics. ChemBioChem 20, 1210–1224 (2019).

12. A. Thompson et al., Tandem mass tags: a novel quantification strategy for comparative analysis of complex protein mixtures by MS/MS. Anal. Chem. 75, 1895–1904 (2003).

13. M. J. Ellis et al., Connecting genomic alterations to cancer biology with proteomics: the NCI Clinical Proteomic Tumor Analysis Consortium. Cancer Discov. 3, 1108–1112 (2013).

14. L. C. Gillet et al., Targeted data extraction of the MS/MS spectra generated by data-independent acquisition: a new concept for consistent and accurate proteome analysis. Mol Cell Proteomics 11, O111.016717 (2012).

15. Y. Perez-Riverol et al., The PRIDE database and related tools and resources in 2019: improving support for quantification data. Nucleic Acids Res. 47, D442–D450 (2019).

16. M. J. Alvarez et al., Functional characterization of somatic mutations in cancer using network-based inference of protein activity. Nat. Genet. 48, 838–847 (2016).

17. M. Wilhelm et al., Mass-spectrometry-based draft of the human proteome. Nature 509, 582–587 (2014).

18. N. Fortelny, C. M. Overall, P. Pavlidis, G. V. C. Freue, Can we predict protein from mRNA levels? Nature 547, E19–E20 (2017).

19. C. Buccitelli, M. Selbach, mRNAs, proteins and the emerging principles of gene expression control. Nat. Rev. Genet. 21, 630–644 (2020).

20. S. Tasaki et al., Inferring protein expression changes from mRNA in Alzheimer’s dementia using deep neural networks. Nat Commun 13, 655 (2022).

21. J. C. Taggart, H. Zauber, M. Selbach, G. W. Li, E. McShane, Keeping the Proportions of Protein Complex Components in Check. Cell Syst 10, 125–132 (2020).

22. A. L. Richards, M. Eckhardt, N. J. Krogan, Mass spectrometry-based protein-protein interaction networks for the study of human diseases. Mol. Syst. Biol. 17, e8792 (2021).

23. M. Giurgiu et al., CORUM: the comprehensive resource of mammalian protein complexes-2019. Nucleic Acids Res. 47, D559–D563 (2019).

24. A. B. Lopacinski et al., Modeling the complete kinetics of coxsackievirus B3 reveals human determinants of host-cell feedback. Cell Syst 12, 304–323 e313 (2021).

25. C. D. Griffiths, A. J. Sweatt, K. A. Janes, Simulating coxsackievirus B3 infection with an accessible computational model of its complete kinetics. STAR Protoc 2, 100940 (2021).

26. G. Ciriello et al., Comprehensive Molecular Portraits of Invasive Lobular Breast Cancer. Cell 163, 506–519 (2015).

27. D. P. Nusinow et al., Quantitative Proteomics of the Cancer Cell Line Encyclopedia. Cell 180, 387–402 e316 (2020).

28. Y. Liu et al., Impact of Alternative Splicing on the Human Proteome. Cell Rep 20, 1229–1241 (2017).

29. G. Rosenberger et al., A repository of assays to quantify 10,000 human proteins by SWATH-MS. Sci Data 1, 140031 (2014).

30. H. Srivastava et al., Protein prediction models support widespread post-transcriptional regulation of protein abundance by interacting partners. PLoS Comput. Biol. 18, e1010702 (2022).

31. M. Ghandi et al., Next-generation characterization of the Cancer Cell Line Encyclopedia. Nature 569, 503–508 (2019).

32. D. G. Mun et al., Proteogenomic Characterization of Human Early-Onset Gastric Cancer. Cancer Cell 35, 111–124 e110 (2019).

33. A. Ruepp et al., CORUM: the comprehensive resource of mammalian protein complexes--2009. Nucleic Acids Res. 38, D497–501 (2010).

34. E. J. Wagenmakers, S. Farrell, AIC model selection using Akaike weights. Psychon Bull Rev 11, 192–196 (2004).

35. M. Wuhr et al., Deep proteomics of the Xenopus laevis egg using an mRNA-derived reference database. Curr. Biol. 24, 1467–1475 (2014).

36. D. Szklarczyk et al., The STRING database in 2023: protein-protein association networks and functional enrichment analyses for any sequenced genome of interest. Nucleic Acids Res. 51, D638–D646 (2023).

37. Y. Liu et al., Multi-omic measurements of heterogeneity in HeLa cells across laboratories. Nat. Biotechnol. 37, 314–322 (2019).

38. J. Reimegard et al., A combined approach for single-cell mRNA and intracellular protein expression analysis. Commun Biol 4, 624 (2021).

39. A. D. Brunner et al., Ultra-high sensitivity mass spectrometry quantifies single-cell proteome changes upon perturbation. Mol. Syst. Biol. 18, e10798 (2022).

40. M. Wang, C. J. Herrmann, M. Simonovic, D. Szklarczyk, C. von Mering, Version 4.0 of PaxDb: Protein abundance data, integrated across model organisms, tissues, and cell-lines. Proteomics 15, 3163–3168 (2015).

41. B. Eraslan et al., Quantification and discovery of sequence determinants of protein-per-mRNA amount in 29 human tissues. Mol. Syst. Biol. 15, e8513 (2019).

42. W. C. Reinhold et al., RNA Sequencing of the NCI-60: Integration into CellMiner and CellMiner CDB. Cancer Res. 79, 3514–3524 (2019).

43. T. Guo et al., Quantitative Proteome Landscape of the NCI-60 Cancer Cell Lines. iScience 21, 664–680 (2019).

44. K. Charmpi et al., Convergent network effects along the axis of gene expression during prostate cancer progression. Genome Biol. 21, 302 (2020).

45. S. Y. Ow et al., iTRAQ underestimation in simple and complex mixtures: “the good, the bad and the ugly”. J. Proteome Res. 8, 5347–5355 (2009).

46. K. A. Janes, An analysis of critical factors for quantitative immunoblotting. Sci Signal 8, rs2 (2015).

47. K. S. Kim, G. Hufnagel, N. M. Chapman, S. Tracy, The group B coxsackieviruses and myocarditis. Rev. Med. Virol. 11, 355–368 (2001).

48. G. Consortium, The GTEx Consortium atlas of genetic regulatory effects across human tissues. Science 369, 1318–1330 (2020).

49. C. F. Liu et al., Whole-Transcriptome Profiling of Human Heart Tissues Reveals the Potential Novel Players and Regulatory Networks in Different Cardiomyopathy Subtypes of Heart Failure. Circ Genom Precis Med 14, e003142 (2021).

50. M. Heinig et al., Natural genetic variation of the cardiac transcriptome in non-diseased donors and patients with dilated cardiomyopathy. Genome Biol. 18, 170 (2017).

51. H. Fechner et al., Induction of coxsackievirus-adenovirus-receptor expression during myocardial tissue formation and remodeling: identification of a cell-to-cell contact-dependent regulatory mechanism. Circulation 107, 876–882 (2003).

52. T. Kaur et al., Expression of coxsackievirus and adenovirus receptor and its cellular localization in myocardial tissues of dilated cardiomyopathy. Exp. Clin. Cardiol. 17, 183–186 (2012).

53. T. H. Dunnebacke, M. B. Reaume, Correlation of the yield of poliovirus with the size of isolated tissue cultured cells. Virology 6, 8–13 (1958).

54. K. A. Hoadley et al., Cell-of-Origin Patterns Dominate the Molecular Classification of 10,000 Tumors from 33 Types of Cancer. Cell 173, 291–304 e296 (2018).

55. R. A. Neff et al., Molecular subtyping of Alzheimer’s disease using RNA sequencing data reveals novel mechanisms and targets. Sci Adv 7, (2021).

56. R. O. Ramirez Flores et al., Consensus Transcriptional Landscape of Human End-Stage Heart Failure. J Am Heart Assoc 10, e019667 (2021).

57. C. M. Perou et al., Molecular portraits of human breast tumours. Nature 406, 747–752 (2000).

58. J. S. Parker et al., Supervised risk predictor of breast cancer based on intrinsic subtypes. J. Clin. Oncol. 27, 1160–1167 (2009).

59. N. Cancer Genome Atlas, Comprehensive molecular portraits of human breast tumours. Nature 490, 61–70 (2012).

60. R. S. Finn et al., Palbociclib and Letrozole in Advanced Breast Cancer. N. Engl. J. Med. 375, 1925–1936 (2016).

61. D. J. Slamon et al., Overall Survival with Ribociclib plus Fulvestrant in Advanced Breast Cancer. N. Engl. J. Med. 382, 514–524 (2020).

62. R. S. Finn et al., Biomarker Analyses of Response to Cyclin-Dependent Kinase 4/6 Inhibition and Endocrine Therapy in Women with Treatment-Naive Metastatic Breast Cancer. Clin Cancer Res 26, 110–121 (2020).

63. N. C. Turner et al., Cyclin E1 Expression and Palbociclib Efficacy in Previously Treated Hormone Receptor-Positive Metastatic Breast Cancer. J. Clin. Oncol. 37, 1169–1178 (2019).

64. C. Vogel et al., Sequence signatures and mRNA concentration can explain two-thirds of protein abundance variation in a human cell line. Mol. Syst. Biol. 6, 400 (2010).

65. B. Schwanhausser et al., Global quantification of mammalian gene expression control. Nature 473, 337–342 (2011).

66. R. Zhang, N. F. Lahens, H. I. Ballance, M. E. Hughes, J. B. Hogenesch, A circadian gene expression atlas in mammals: implications for biology and medicine. Proc. Natl. Acad. Sci. U. S. A. 111, 16219–16224 (2014).

67. J. Lever, M. Krzywinski, N. Altman, Regularization. Nat. Methods 13, 803–804 (2016).

68. E. Schneidman, M. J. Berry, 2nd, R. Segev, W. Bialek, Weak pairwise correlations imply strongly correlated network states in a neural population. Nature 440, 1007–1012 (2006).

69. H. Haken, Information compression in biological systems. Biol. Cybern. 56, 11–17 (1987).

70. E. Brunk et al., Recon3D enables a three-dimensional view of gene variation in human metabolism. Nat. Biotechnol. 36, 272–281 (2018).

71. S. A. Becker, B. O. Palsson, Context-specific metabolic networks are consistent with experiments. PLoS Comput. Biol. 4, e1000082 (2008).

72. R. Agren et al., Reconstruction of genome-scale active metabolic networks for 69 human cell types and 16 cancer types using INIT. PLoS Comput. Biol. 8, e1002518 (2012).

73. S. Opdam et al., A Systematic Evaluation of Methods for Tailoring Genome-Scale Metabolic Models. Cell Syst 4, 318–329 e316 (2017).

74. C. Q. Zhong et al., Generation of a murine SWATH-MS spectral library to quantify more than 11,000 proteins. Sci Data 7, 104 (2020).

75. B. Efron, R. Tibshirani, Statistical data analysis in the computer age. Science 253, 390–395 (1991).

76. R. Gao et al., Delineating copy number and clonal substructure in human tumors from single-cell transcriptomes. Nat. Biotechnol. (2021).

77. W. Saelens, R. Cannoodt, H. Todorov, Y. Saeys, A comparison of single-cell trajectory inference methods. Nat. Biotechnol. 37, 547–554 (2019).

78. M. C. Chambers et al., A cross-platform toolkit for mass spectrometry and proteomics. Nat. Biotechnol. 30, 918–920 (2012).

79. H. L. Rost et al., OpenMS: a flexible open-source software platform for mass spectrometry data analysis. Nat. Methods 13, 741–748 (2016).

80. J. Teleman et al., DIANA--algorithmic improvements for analysis of data-independent acquisition MS data. Bioinformatics 31, 555–562 (2015).

81. H. L. Rost et al., TRIC: an automated alignment strategy for reproducible protein quantification in targeted proteomics. Nat. Methods 13, 777–783 (2016).

82. R. Leinonen, H. Sugawara, M. Shumway, C. International Nucleotide Sequence Database, The sequence read archive. Nucleic Acids Res. 39, D19–21 (2011).

83. D. Kim, J. M. Paggi, C. Park, C. Bennett, S. L. Salzberg, Graph-based genome alignment and genotyping with HISAT2 and HISAT-genotype. Nat. Biotechnol. 37, 907–915 (2019).

84. H. Li et al., The Sequence Alignment/Map format and SAMtools. Bioinformatics 25, 2078–2079 (2009).

85. S. Kovaka et al., Transcriptome assembly from long-read RNA-seq alignments with StringTie2. Genome Biol. 20, 278 (2019).

86. G. H. Putri, S. Anders, P. T. Pyl, J. E. Pimanda, F. Zanini, Analysing high-throughput sequencing data in Python with HTSeq 2.0. Bioinformatics 38, 2943–2945 (2022).

87. J. Morales et al., A joint NCBI and EMBL-EBI transcript set for clinical genomics and research. Nature 604, 310–315 (2022).

88. D. P. Nusinow, S. P. Gygi, A Guide to the Quantitative Proteomic Profiles of the Cancer Cell Line Encyclopedia. bioRxiv 2020.2002.2003.932384 (2020).

89. C. Gene Ontology et al., The Gene Ontology knowledgebase in 2023. Genetics 224, (2023).

90. W. Shao et al., Comparative analysis of mRNA and protein degradation in prostate tissues indicates high stability of proteins. Nat Commun 10, 2524 (2019).

91. C. R. John et al., M3C: Monte Carlo reference-based consensus clustering. Sci. Rep. 10, 1816 (2020).

