## Supplemental Figures and Methods for "Proteome-wide copy-number estimation from transcriptomics"

### SI Appendix, Materials and Methods

**Median (M) regression.** For each gene, the median was calculated across the SWATH-scaled proteomic dataset, taking empty entries as missing elements rather than zeros. A 95% confidence interval of the median was estimated by bootstrapping (*n* = 1000 runs).

**Hyperbolic-to-linear (HL) regression.** For each gene, an HL model was constructed as follows:

$$SWATH\text{–}\ scaled\ protein\ copies\ per\ cell\ =\ a \bullet \left(\frac{b \bullet TPM}{c+TPM} + TPM\right),$$

where *a*, *b*, and *c* were regression coefficients estimated by nonlinear least squares in MATLAB (version R2022a). To prevent discontinuities from division by zero, *c* was constrained to be greater than zero. Additionally, a logistic weighting of the cost function was desired to prevent high-abundance cell lines from overleveraging the regression. To achieve both, we devised a two-step procedure in which initial estimates were made with a linear cost function and $c > 0$ constraint using lsqcurvefit with 'FiniteDifferenceType' set to 'Central'. The regression estimates of *a*, *b*, and *c* were then used as an initial guess for nlinfit with 'RobustWgtFun' set to 'logistic', and the regression was repeated without constraints. If the updated value of c was less than zero, then the updated regression estimates of *a* and *b* were used as initial guesses for a second round of lsqcurvefit with a linear cost function and $c > 0$ constraint. A 95% confidence interval of the HL fit was estimated by asymptotic error analysis of the regression coefficients using the *F* distribution to describe the ratio of the sum-of-squared errors for the ideal and parameter-perturbed model divided by their corresponding degrees of freedom.

**HL + least absolute shrinkage and selection operator (HL+LASSO) regression.** Residuals from the HL fit were regressed against all other genes in the transcriptome by LASSO with the glmnet package (version 4.1-6) in R. To determine the optimal penalty strength parameter ($\lambda$) for each LASSO regression, we used the cv.glmnet function for cross-validation after increasing the function's minimum fractional change in deviance for stopping (fdev) to 0.01. We accounted for differences among cross-validation runs by iterating cv.glmnet 100 times, calculating the BIC for the best $\lambda$ in each iteration, and defining the best $\lambda$ with the lowest BIC as optimal. The optimal $\lambda$ was used with the glmnet function and all observations of the gene to obtain a regularized feature set and linear coefficients. Output of the LASSO regression was subtracted from HL fit to obtain the HL+LASSO model. LASSO displacements were propagated linearly to the 95% confidence interval of the HL fit, and graphical displays were LOESS smoothed with the geom_smooth function in ggplot2 (version 3.4.0) in R.

**Comparison of HL model to alternatives.** For each gene, three alternative models were constructed as follows:

Linear: $SWATH\text{–}\ scaled\ protein\ copies\ per\ cell\ =\ a \bullet TPM$

Hyperbolic: $SWATH\text{–}\ scaled\ protein\ copies\ per\ cell\ = \frac{a \bullet TPM}{b+TPM};\ b > 0$

3-parameter logistic: $SWATH\text{–}\ scaled\ protein\ copies\ per\ cell\ = a - \frac{a}{1+(\frac{TPM}{b})^c};\ a > 0, b > 0$

where *a*, *b*, and *c* were regression coefficients to be estimated in MATLAB (version R2022a). Linear models were regressed using fitlm with 'RobustOpts' set to 'logistic' and 'Intercept' set to false. Hyperbolic and 3-parameter logistic models were fit similarly to HL but with a combination of lsqcurvefit for constrained regression and fitnlm (which calls nlinfit internally) for logistic weighting of the cost function and the estimation of log-likelihood, which was used for Bayesian Information Criterion (BIC) calculation with the aicbic function. BIC weights (BICw) were calculated as follows:

$$BICw_i\ = \frac{exp\left[-\frac{1}{2}(BIC_i - BIC_{min})\right]}{\sum_{k=1}^{K} exp\left[-\frac{1}{2}(BIC_k - BIC_{min})\right]}$$

where $BIC_i$ is the BIC for the $i^{th}$ model and $BIC_{min}$ is the minimum BIC in the group of models (1).

**STRING visualization.** Interaction maps were drafted on the STRING database web site (2) under Search>Multiple proteins. The default output maps were altered as follows: meaning of network edges

= confidence, minimum required interaction score = medium confidence (0.400), disable 3D bubble design, and disable structure previews inside network bubbles. Interaction maps were exported as vectorized SVG files for further stylistic refinement.

**HeLa RNA-seq downsampling.** Before downsampling, RNA-seq data from the HeLa derivatives were averaged. Raw counts were converted to counts per million (CPM), and the counts per million-normalized reads for each gene were averaged across all derivatives. Then, the average CPM was converted back to an averaged count by multiplying the average read depth and rounding to the nearest integer. The average counts for each gene were downsampled 100 times using rbinom in R, with the number of trials equal to the downsampled read depth (25 million to 50,000) and the probability of success equal to the number of average counts for that gene divided by the total number of average counts. Downsampled counts were converted to TPM and used with Pinferna to predict the mean-averaged SWATH data from the HeLa derivatives.

**Hierarchical clustering.** Breast cancer RNA-seq was $\log_2$-transformed and row-standardized. For inferred proteomic profiles, M genes were removed before standardization because of zero variance. Data were clustered by Euclidean distance with Ward's linkage using the Heatmap function in the ComplexHeatmap (version 2.12.1) package. Columns were clustered within a subtype defined either by transcriptomics or proteomics. To identify genes that changed disproportionately between mRNA and protein, the Z-scores of the species were subtracted: $Z_{diff}$ = Z-score$_{Protein}$ - Z-score$_{mRNA}$. To identify genes of interest that drove luminal reassignments, we filtered for genes with a $|Z_{diff}| \geq 3$ and ranked by the frequency of occurrence, focusing on genes with $|Z_{diff}| \geq 3$ in six or more samples that had undergone subtype reassignment.

### SI Appendix, Figures

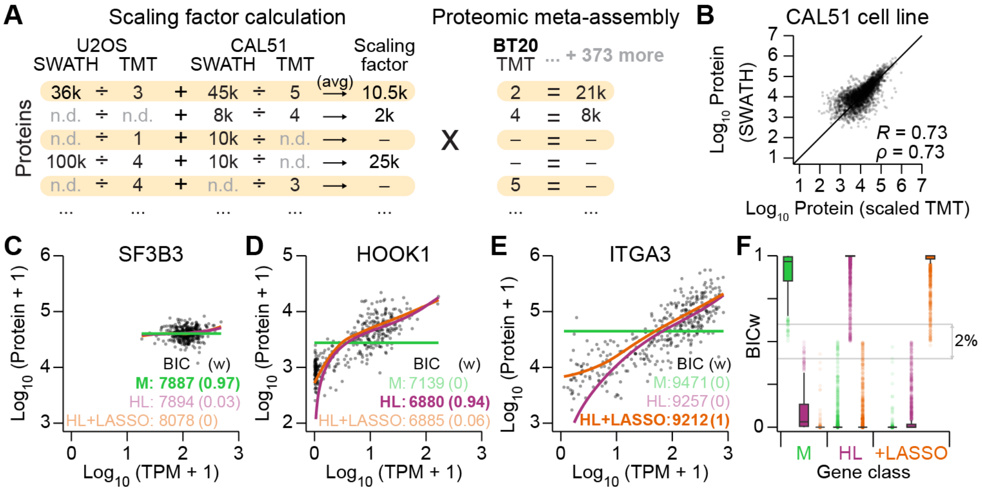


**Fig. S1.** Calibration for the proteomic meta-assembly and model selection examples.

(*A*) SWATH–TMT scaling factor estimation and calibration of the proteomic meta-assembly. Scaling factors were estimated for each protein by dividing the SWATH intensity by the TMT intensity in U2OS and CAL51 when both data types were available and averaging (avg) the two ratios when possible. The gene-specific scaling factors were used to convert the entire TMT dataset (bold) to absolute protein abundances.

(*B*) The reciprocal cross-calibration to that shown in Fig. 1*B*. Step 1 of Fig. 1*A* was performed with U2OS data alone and the SWATH-scaled TMT proteomics of CAL51 cells compared with data obtained directly by SWATH. Pearson's *R* and Spearman's *ρ* are shown.

(*C–E*) Model selection of the representative genes shown in Fig. 1*C–E*. Absolute protein copies per cell were regressed against the mRNA abundance normalized as transcripts per million (TPM) for $n$ = 369 cancer cell lines. Data are fit with M, HL, and HL+LASSO models. The BIC was used to discriminate the best model for each fit, and BIC weights (w) are shown in parentheses to indicate the relative best-model probability. The lowest BIC is indicated.

(*F*) BIC weights (BICw) for each model fit to M, HL, or HL+LASSO (+LASSO) genes. Range of weights for genes with model ambiguity (two models with a BICw ≥ 0.4) are boxed (gray) with the percentage indicated.

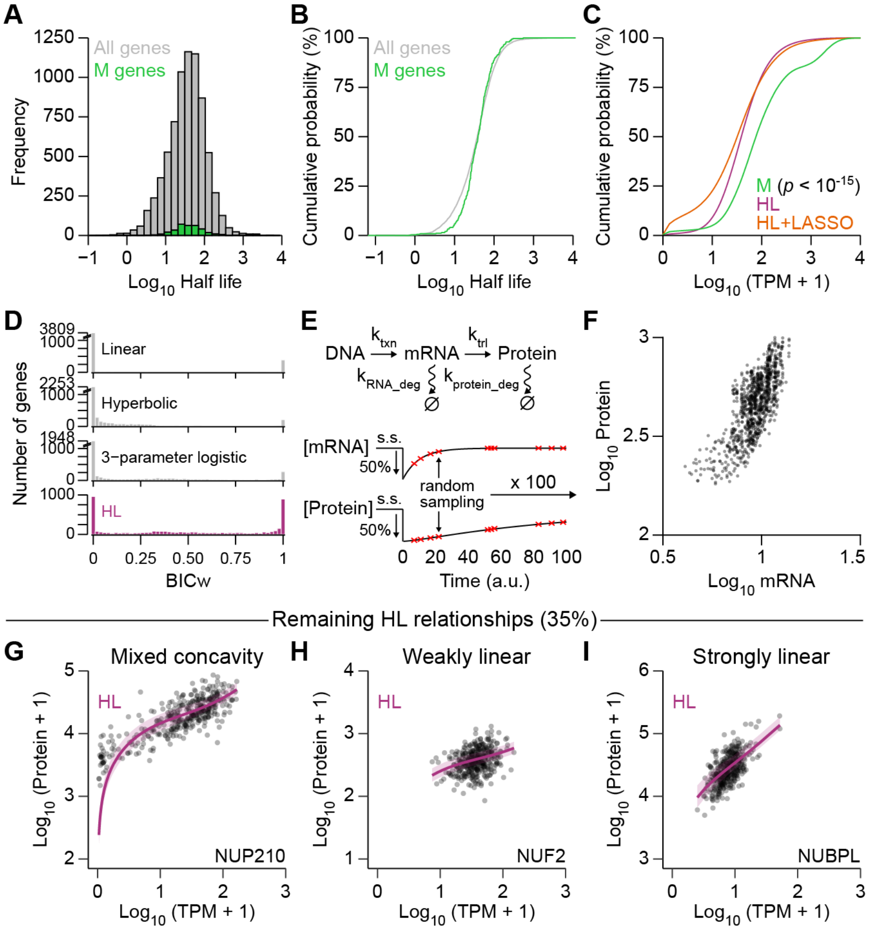


**Fig. S2.** Extended characterization of M and HL mRNA-to-protein relationship classes.
(*A, B*) M genes do not exhibit longer half-lives compared to other relationship classes. Half-lives for proteins were obtained from (3) and plotted by frequency (*A*) or as an empirical cumulative distribution function (*B*) for all genes ($n$ = 7029 genes (3); gray) and M genes ($n$ = 334 genes; green).
(*C*) M genes are more abundant by mRNA compared to other relationship classes. mRNA abundance was normalized as TPM and placed on a log scale with a pseudocount of 1. Cumulative distribution functions are shown for M genes ($n$ = 395 genes; green), HL genes ($n$ = 2569 genes; purple), and HL+LASSO genes ($n$ = 1402 genes; orange). Distributions were compared by K-S test with Šidák correction for multiple-hypothesis testing.
(*D*) Distribution of BIC weights (BICw) for models encoding linear, hyperbolic, three-parameter logistic, and HL relationships ($n$ = 4366 genes) shown in Fig. 2*B*.
(*E*, *F*) Log concave-up patterns arise when mRNA and protein abundances recover from transient perturbations to steady-state values. A simple transcription–translation model (*E*) was reduced by 50% and randomly sampled at 10 time points (red) during the return to steady state (s.s.). For the model, the following dimensionless rate parameters were used: $k_{txn}$ = 1; $k_{trl}$ = 1; $k_{RNA\_deg}$ = 0.1; $k_{protein\_deg}$ = 0.01. The model was simulated 100 times with lognormally distributed parameter noise (coefficient of variation = 10%) and the joint observation of mRNA and protein abundances ($n$ = 10 time points x 100 simulations) is shown in *F*.
(*G*–*I*) Examples of other HL relationships not shown in Fig. 2*C,D*: mixed concavity (*G*), weakly linear (*H*), and strongly linear (*I*) relationships.

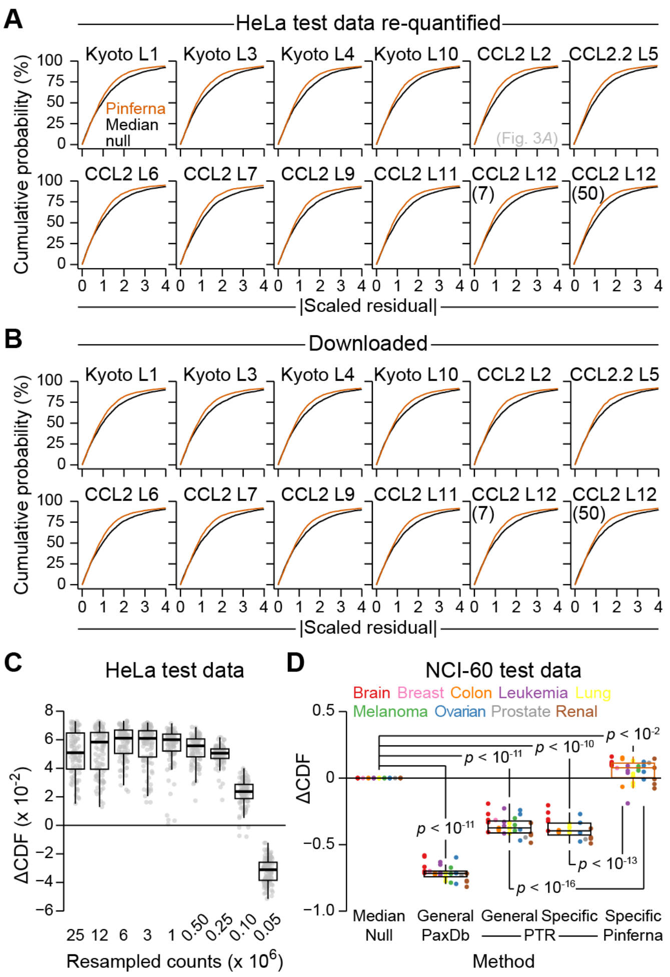


**Fig. S3.** Robustness of Pinferna predictions.

(*A*, *B*) Prediction accuracy is not dependent on SWATH analytical details. Cumulative distribution plots comparing Pinferna and median-null predictions for paired RNA-seq–SWATH datasets of 12 HeLa derivatives (PRJNA437150; PXD009273) named as in the publication (4). Proteins were re-quantified from raw SWATH data processed exactly like the meta-assembly (*Materials and Methods*; *A*); or, protein quantities were taken directly from the publication (4) (*B*). Results from the re-quantified CCL2 L2 derivative are reprinted from Fig. 3*A*.

(*C*) Prediction accuracy is not heavily dependent on RNA-seq read depth. Count-based RNA-seq data for the HeLa lines was averaged and iteratively downsampled ($n$ = 100 iterations; gray) and TPM values re-estimated before making proteome-wide copy-number predictions by Pinferna. See Fig. 3*B* for an explanation of ΔCDF.

(*D*) Prediction accuracy is not dependent on RNA-seq analytical details. Pinferna predictions were made using TPM values taken directly from the original publication (5), and ΔCDF values were calculated for NCI-60 cell lines excluded from model training (Fig. 1*A*) and organized by cancer type ($n$ = 5 brain, 1 breast, 3 colon, 4 leukemia, 4 lung, 3 melanoma, 3 ovarian, 1 prostate, 5 renal). Differences between groups were assessed by rank-sum test with Šidák correction.

For *C* and *D*, box-and-whisker plots show the median (horizontal line), interquartile range (IQR; box), and an additional 1.5 IQR extension (whiskers) of the data.

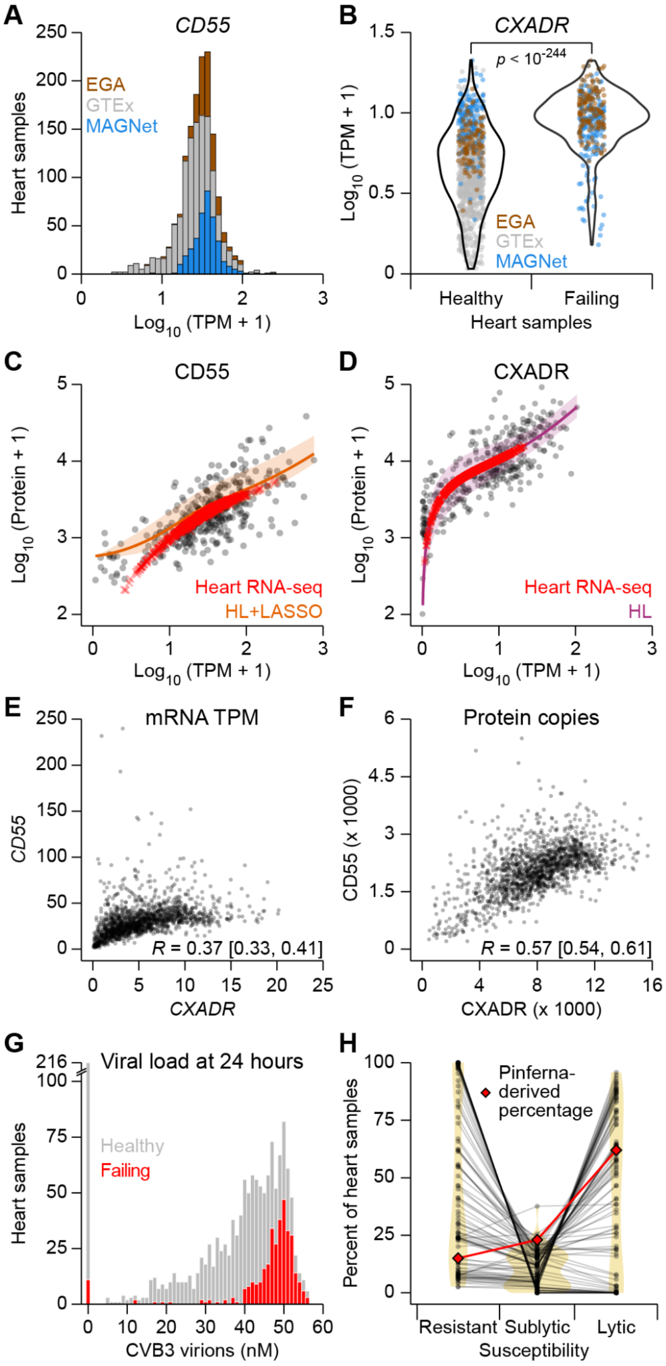


**Fig. S4.** Calibrated protein inferences of CD55 and CXADR yield disease-related predictions of coxsackievirus B3 (CVB3) susceptibility.

(*A*) Distribution of *CD55* abundance (*SI Appendix*, Table S7) separated by source data: EGA (brown; EGAS00001002454), GTEx (gray; phs000424.v9.p2), MAGNet (blue; GSE141910).

(*B*) *CXADR* is upregulated in failing hearts. *CXADR* abundance (*SI Appendix*, Table S7) was stratified by heart health and colored by source data as in *A*. Differences between groups were assessed by rank-sum test.

(*C, D*) Calibration plots (*C*, orange; *D*, purple) and Pinferna predictions for CD55 (*C*) and CXADR (*D*) in human heart samples (Heart RNA-seq; red). Best-fit calibrations ± 95% confidence intervals are overlaid on the proteomic–transcriptomic data from $n$ = 369 cancer cell lines. CD55 deviations from the smoothed best fit are caused by cardiac-specific features in the HL+LASSO regressions.

(*E, F*) CD55–CXADR coregulation is increased at the protein level (*F*) compared to the mRNA level (*E*). For each comparison, the Pearson *R* is shown with 95% confidence interval in brackets calculated by the Fisher Z transformation.

(*G*) Replotted histogram of Fig. 4*C* separated by heart health (*SI Appendix*, Table S7).

(*H*) Predicted prevalence of susceptibility groups based on randomized measurements. After linearly scaling to randomized abundances of CD55 and CXADR (*n* = 100 randomizations), CVB3 infections were simulated for 1489 heart samples as in Fig. 4*B*. The 24-hour end states were quantified by the percentage of samples with resistant ([CVB3 virions] = 0 nM), sublytic (0 nM < [CVB3 virions] < 36 nM), and lytic ([CVB3 virions] ≥ 36 nM) phenotypes (Fig. 4*B*). Pinferna-derived percentages are overlaid in red. Results from the randomized simulations are connected, and densities in each group are shown by a violin plot in the background (yellow).

For *B*–*F*, *n* = 1489 heart samples.

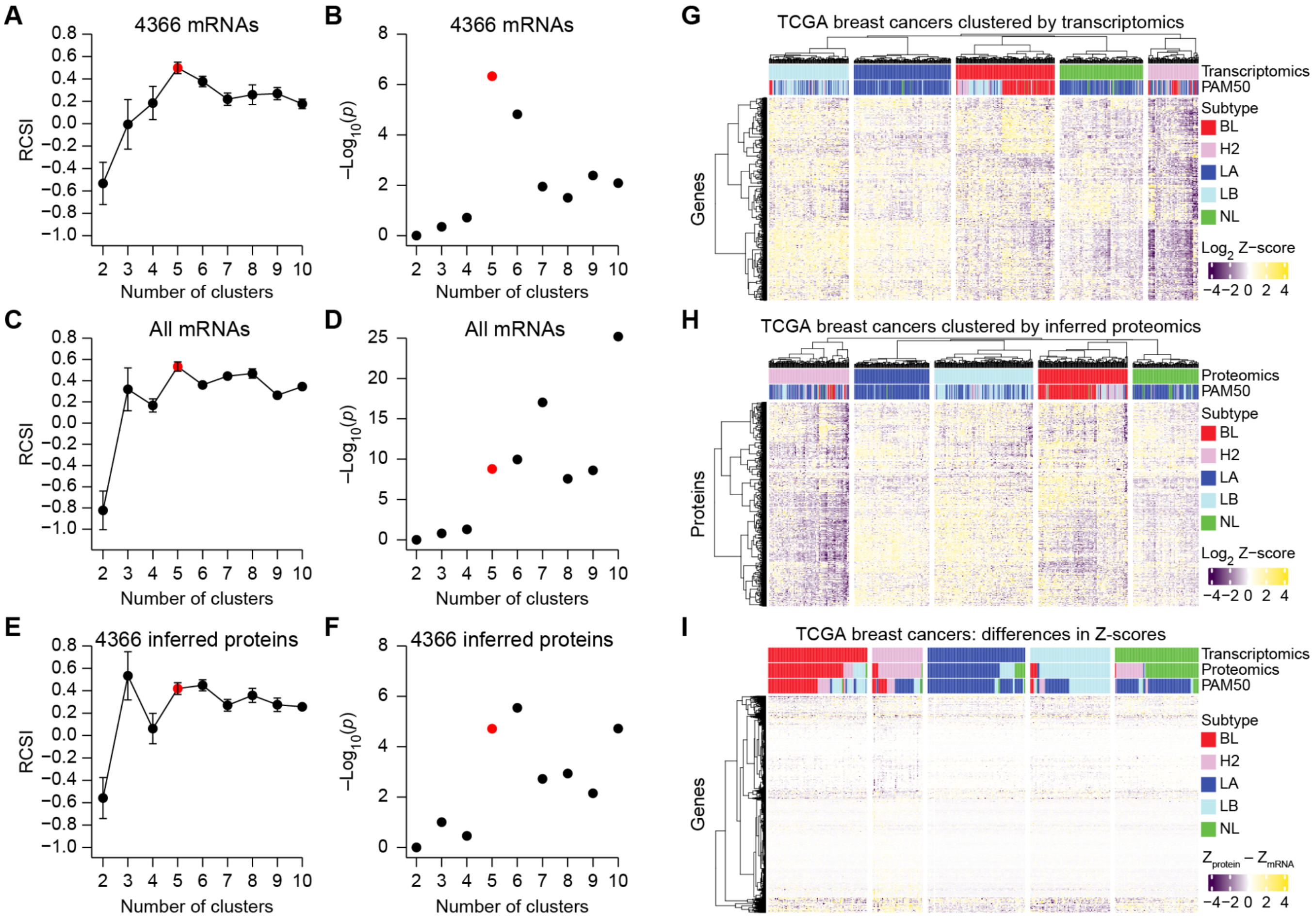


**Fig. S5.** Consensus re-clustering of 796 breast cancer cases from The Cancer Genome Atlas (TCGA) by protein inference.
(*A*–*F*) Relative cluster stability index (RCSI; *A*, *C*, and *E*) and significance (*B*, *D*, and *F*) of cluster number for consensus clustering (*Materials and Methods*) applied to 4366 mRNAs used by Pinferna (*A*, *B*), all mRNAs quantified by RNA-seq (*C*, *D*), and protein inferences by Pinferna (*E*, *F*). Stable maxima or near maxima at five clusters is indicated in red.
(*G*, *H*) Two-way hierarchical clustering within transcriptomic (*G*) and inferred proteomic (*H*) clusters annotated as in Fig. 5*A*.
(*I*) One-way hierarchical clustering of genes by differences in standardized Z-scores between inferred protein and mRNA ($Z_{protein} - Z_{mRNA}$).
For *G*–*I*, hierarchical clustering was performed by Euclidean distance and Ward's linkage. BL: Basal-like; H2: HER2+; LA: Luminal A; LB: Luminal B; NL: Normal-like.

**Supplementary Tables**

**Table S1.** SWATH-scaled proteomics data.
**Table S2.** Harmonized identifier table for 19,062 human genes.
**Table S3.** Harmonized and filtered RNA-seq data used for regression.
**Table S4.** Harmonized and filtered SWATH-scaled proteomics data used for regression.
**Table S5.** HL parameters, LASSO features + coefficients, BICs.
**Table S6.** GO enrichments for M, HL, and HL+LASSO genes.
**Table S7.** Harmonized prostate table.
**Table S8.** CD55 and CXADR TPM together with identifier, study name, and heart status.
**Table S9.** Data template for Pinferna web-based tool.

**SI Appendix, References**


1. E. J. Wagenmakers, S. Farrell, AIC model selection using Akaike weights. *Psychon Bull Rev* **11,** 192-196 (2004).
2. D. Szklarczyk *et al.*, The STRING database in 2023: protein-protein association networks and functional enrichment analyses for any sequenced genome of interest. *Nucleic Acids Res.* **51,** D638-D646 (2023).
3. J. Zecha *et al.*, Peptide Level Turnover Measurements Enable the Study of Proteoform Dynamics. *Mol Cell Proteomics* **17,** 974-992 (2018).
4. Y. Liu *et al.*, Multi-omic measurements of heterogeneity in HeLa cells across laboratories. *Nat. Biotechnol.* **37,** 314-322 (2019).
5. W. C. Reinhold *et al.*, RNA Sequencing of the NCI-60: Integration into CellMiner and CellMiner CDB. *Cancer Res.* **79,** 3514-3524 (2019).
